# Ventral Stress Fibers Induce Plasma Membrane Deformation in Human Fibroblasts

**DOI:** 10.1101/2021.03.01.433420

**Authors:** Samuel J. Ghilardi, Mark S. Aronson, Allyson E. Sgro

**Affiliations:** Department of Biomedical Engineering, Boston University, Boston, MA 02215 USA; Biological Design Center, Boston University, Boston, MA 02215 USA

**Keywords:** actin, stress fibers, plasma membrane, membrane contouring

## Abstract

Interactions between the actin cytoskeleton and the plasma membrane are essential for many eukaryotic cellular processes. During these processes, actin fibers deform the cell membrane outward by applying forces parallel to the fiber’s major axis (as in migration) or they deform the membrane inward by applying forces perpendicular to the fiber’s major axis (as during cytokinesis). Here we describe a novel actin-membrane interaction in human dermal myofibroblasts. When labeled with a cytosolic fluorophore, the myofibroblasts developed prominent fluorescent structures on the ventral side of the cell. These structures are present in the cell membrane and colocalize with ventral actin stress fibers, suggesting that the fibers bend the membrane to form a “cytosolic pocket” for the fluorophores to flow into, creating the observed structures. The existence of this pocket was confirmed by transmission electron microscopy. Dissolving the stress fibers, inhibiting fiber protein binding, or inhibiting myosin II binding of actin removed the observed structures. However, decreasing cellular contractility did not remove the structures. Taken together, our results illustrate a novel actin-membrane bending topology where the membrane is deformed outwards rather than being pinched inwards, resembling the topological inverse of cytokinesis.

## Introduction

Critical cellular processes ranging from contraction (1–3) and migration (1, 4–6) to proliferation (7–11) are dependent on mechanotransduction mediated through actin fibers (5, 12–16). In many cell types, including fibroblasts, actin fibers associate together with myosin proteins to form actin stress fibers (5, 17–19), which provide mechanical integrity and generate contractile forces in the cell. As part of these processes, actin stress fibers interact with the cell’s plasma membrane, most commonly at the end of the fiber through adapter proteins (such as talins or vinculin) at focal adhesions (20–22). These focal adhesion-stress fiber interactions are a common way to classify stress fibers, depending on if the fiber is coupled to focal adhesions on both ends (ventral stress fiber), on one end (dorsal stress fiber), or not coupled to a focal adhesion (transverse arc) (23). In addition to the coupling of one or both ends of stress fibers to focal adhesions, there are also adapter proteins that couple the membrane to the fiber along the length of the stress fiber, such as the ezrin/radixin/moesin (ERM) family of proteins (24–26). Through these protein-membrane interactions, actin stress fibers deform the membrane at smaller structures such as focal adhesions (27, 28) and filopodia, as well as in larger projections like lamellipodia (29–31).

To deform the plasma membrane, actin stress fibers apply force along their principle axis, either through actin polymerization (32–34) or via myosin II contraction (35–38). As a result, during most contractile events, actin stress fibers generally deform the membrane parallel to the major axis of the fiber. A notable exception to this is during specialized membrane pinching events, such as cytokinesis, where actin fibers assemble into a ring-like structure and pull the membrane inward, perpendicular to the axis of the fiber (39–42). In larger structures such as lamellipodia, the arp2/3 complex allows for branching of actin fibers (43, 44), and the generation of complex membrane contours (45–48), but each individual fiber applies forces axially along the fiber. So far, no reported actin structure applies forces perpendicular along the length of the fiber (as in cytokinesis) but deforms the membrane outward (as in filopodia or lamellipodia formation). Here, we report such a structure: an actin-induced membrane bending that occurs along the length of the actin fiber, generating a stable cytosolic pocket. We find this pocket requires direct coupling of the fiber to the membrane but once formed, is independent of active, myosin II-mediated fiber contraction.

## Results and Discussion

### Novel Fluorescent Structures are Observed in Myofibroblasts Labeled with a Cytosolic Fluorophore

After exposure to Transforming Growth Factor Beta (TGF-*β*1), fibroblasts transition to a myofibroblast phenotype (49, 50), characterized by smooth muscle actin *α*-SMA expression (51), as well as an increase in stress fiber formation (50) and cellular contractility (52). We stimulated this transition in human dermal fibroblasts (HDFs) by culturing them with TGF-*β* 1-containing media for 96 hours, after which the HDFs expressed *α*-SMA (Figure S1A) and developed prominent stress fibers, with some *α*-SMA incorporation (Figure S1B). Surprisingly, during this transition, live HDFs loaded with cytosolic fluorescent dye developed fluorescent structures on the ventral side of the cell as seen via spinning disc confocal microscopy (Figure 1A). To test whether these structures were a dye-specific artifact, we generated HDF cell lines expressing the fluorescent proteins mNeonGreen or mScarlet-i, and treated them with TGF-*β*1. Fluorescent structures were observed in both cell lines (Figure 1B&C), suggesting that these observed fluorescent structures are not an artifact of a particular dye. As a side observation, fluorescent puncta were observed in mScarlet-i expressing cells, but not cells expressing mNeongreen. Due to its mCherry lineage (53), it is possible that these puncta are similar to the mCherry aggregations previously observed in mouse neurons (54).

**Fig. 1.**
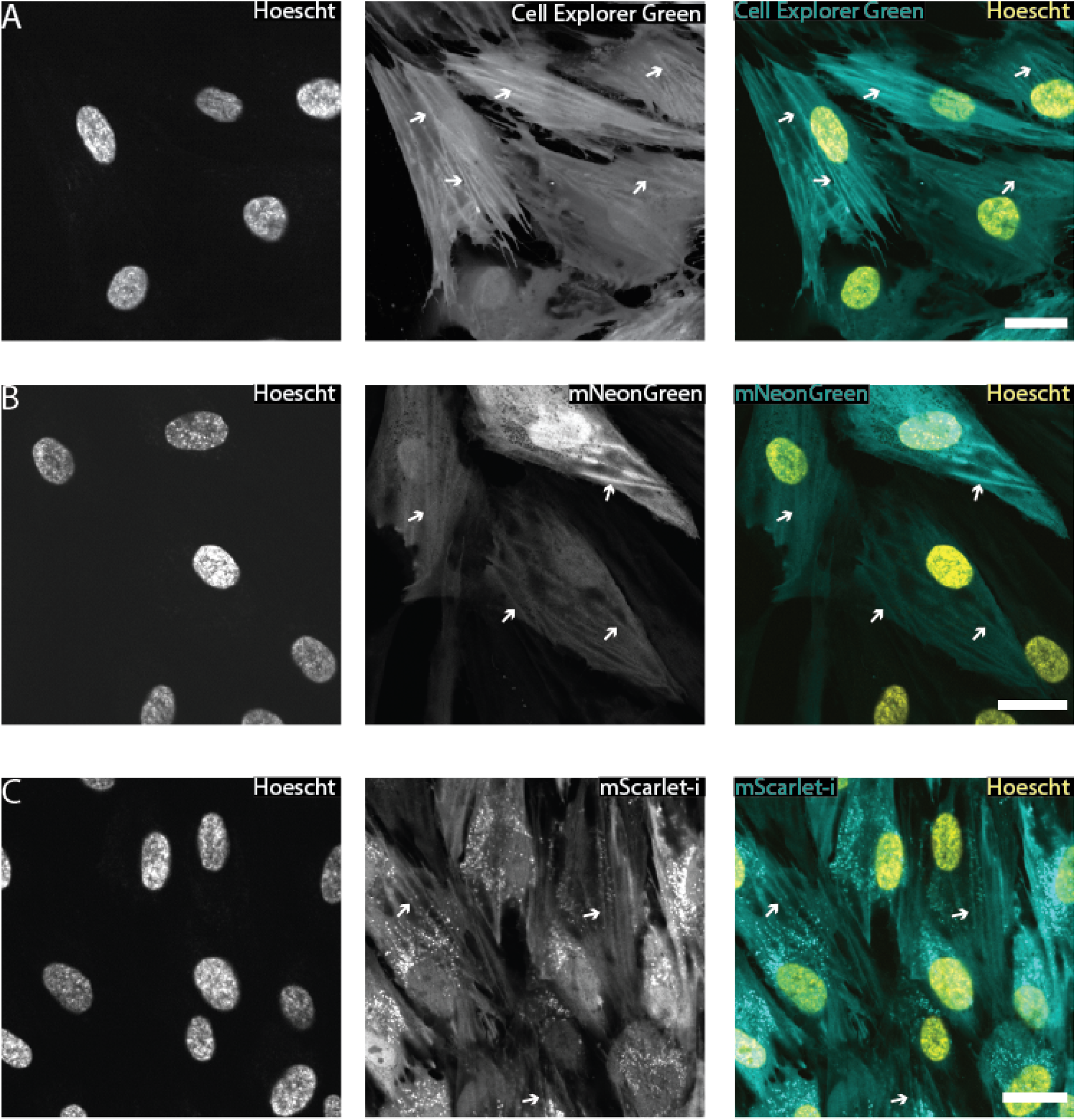
Fluorescent Structures are Visible in Human Dermal Myofibroblasts Loaded with a Cytosolic Fluorophore. After 96 hours of TGF-*β*1 treatment, Human Dermal Fibroblasts transition into myofibroblasts (Figure S1) and separately develop fluorescent structures on the ventral side of the cell (examples marked by white arrows). These ridges can be observed in naive cells labeled with **(A)** cell permeable dye, or cells expressing fluorescent proteins such as **(B)** mNeonGreen or **(C)** mScarlet-i. Note that, at this magnification, fluorescent puncta can be seen in cells expressing either mScarlet-i or mCherry (Figure 5), but not mNeonGreen. There is also some visible bleedthrough from the blue (Hoescht) channel into the green (cell explorer/mNeongreen) channel. Scale Bar = 25 *μ*m. Each experiment was conducted in parallel in three separate wells, and a representative image from one well is shown.

### Observed Fluorescent Structures Colocalize with Ventral Actin Stress Fibers

The observed fluorescent structures appear superficially similar to actin stress fibers, so we stained the cells with phalloidin to observe actin stress fibers. The structures colocalized with actin stress fibers (Figure 2A&C), so we hypothesized that the formation of these structures is related to stress fiber formation. Therefore, we decided to classify the type of stress fiber that colocalized with these ridges.

**Fig. 2.**
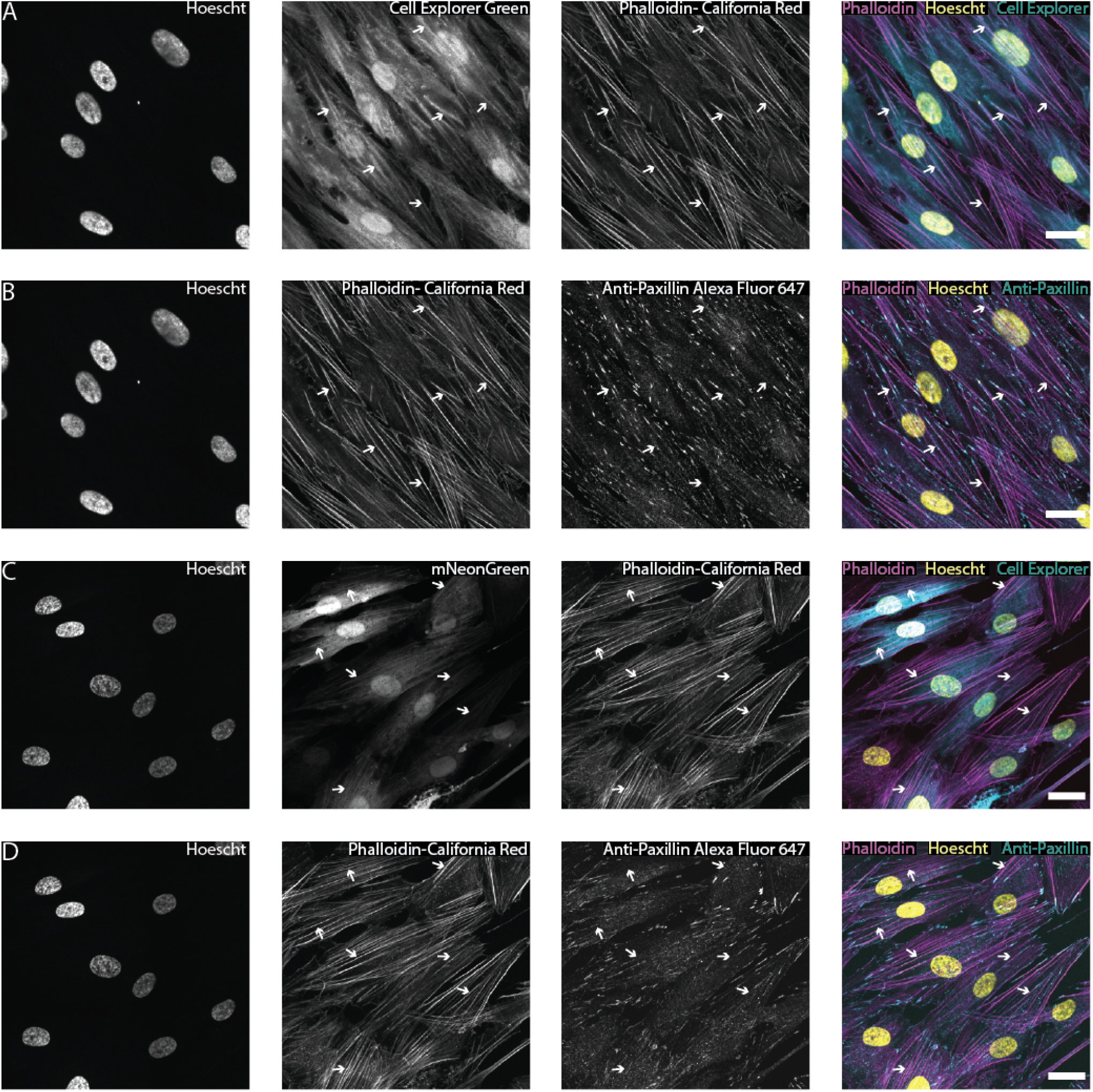
Fluorescent Structures Colocalize with Ventral Actin Stress Fibers. **(A&B)** Cells either stained with green Cell Explorer dye or expressing mNeonGreen **(C&D)** were fixed and stained with Hoescht (nuclei), phalloidin-California Red (actin), and an anti-phospho-paxillin primary antibody (focal adhesions) with an Alexa Fluor-647 secondary. The fluorescent structures (examples marked with white arrows) observed with either the **(A)** Cell Explorer dye or **(C)** mNeonGreen colocalize with phalloidin-stained stress fibers. The colocalized fibers have focal adhesions on both ends of the fiber **(B&D)**, identifying them as ventral stress fibers. Scale Bar = 25 *μ*m A&B and C&D are different channels for the same field of view. Note: there is also some visible bleedthrough from the blue (Hoescht) channel into the green (cell Explorer/mNeongreen) channel. Each experiment was conducted in parallel in three separate wells, and a representative image from one well is shown.

In 2D cell culture, there are three kinds of actin stress fibers: ventral stress fibers, dorsal stress fibers, and transverse arcs (23, 55–58). The different types of fibers can be distinguished by their association with focal adhesions. Specifically, ventral stress fibers are associated with focal adhesions on both ends of the fiber, dorsal stress fibers are associated with a focal adhesion on one end of the fiber, while transverse arcs are not associated with focal adhesions at all. We stained the cells with an anti-phospho-paxillin antibody to observe focal adhesions (Figure 2B&D), which revealed that the majority of colocalized stress fibers started and ended at focal adhesions, identifying them as ventral stress fibers. We also observed that the focal adhesions overlapped with the fluorescent structures (Figure 2B&D), suggesting that focal adhesions, as well as actin stress fibers, could play a role in fluorescent structure formation.

### Fluorescent Structures are Formed by Stress Fiber-Induced Plasma Membrane Deformation Along the Fiber Length

After determining that the observed structures colocalize with ventral actin stress fibers, we investigated potential mechanisms for how the stress fibers cause these structures. One possibility was that the dyes or fluorescent proteins were binding directly to stress fibers. However, the cytosolic Cell Explorer dye, mNeonGreen (59), nor mScarlet-i (53) have any reported intrinsic affinity for actin. Indeed, in the case of the fluorescent proteins, visualization of actin has previously necessitated their fusion to actin binding moieties such as F-tractin or lifeAct (60, 61). Alternatively, the ventral stress fiber could be deforming the plasma membrane, creating a cytosolic “pocket” in the membrane around the stress fiber (Figure 3A). The fluorescent markers then diffuse into this pocket, creating the fluorescent structures observed in Figure 1. As a corollary to this hypothesis, we would expect to see fluorescent structures around focal adhesions, as focal adhesions also cause membrane deformation, as seen via transmission electron microscopy (TEM) (27–29).

**Fig. 3.**
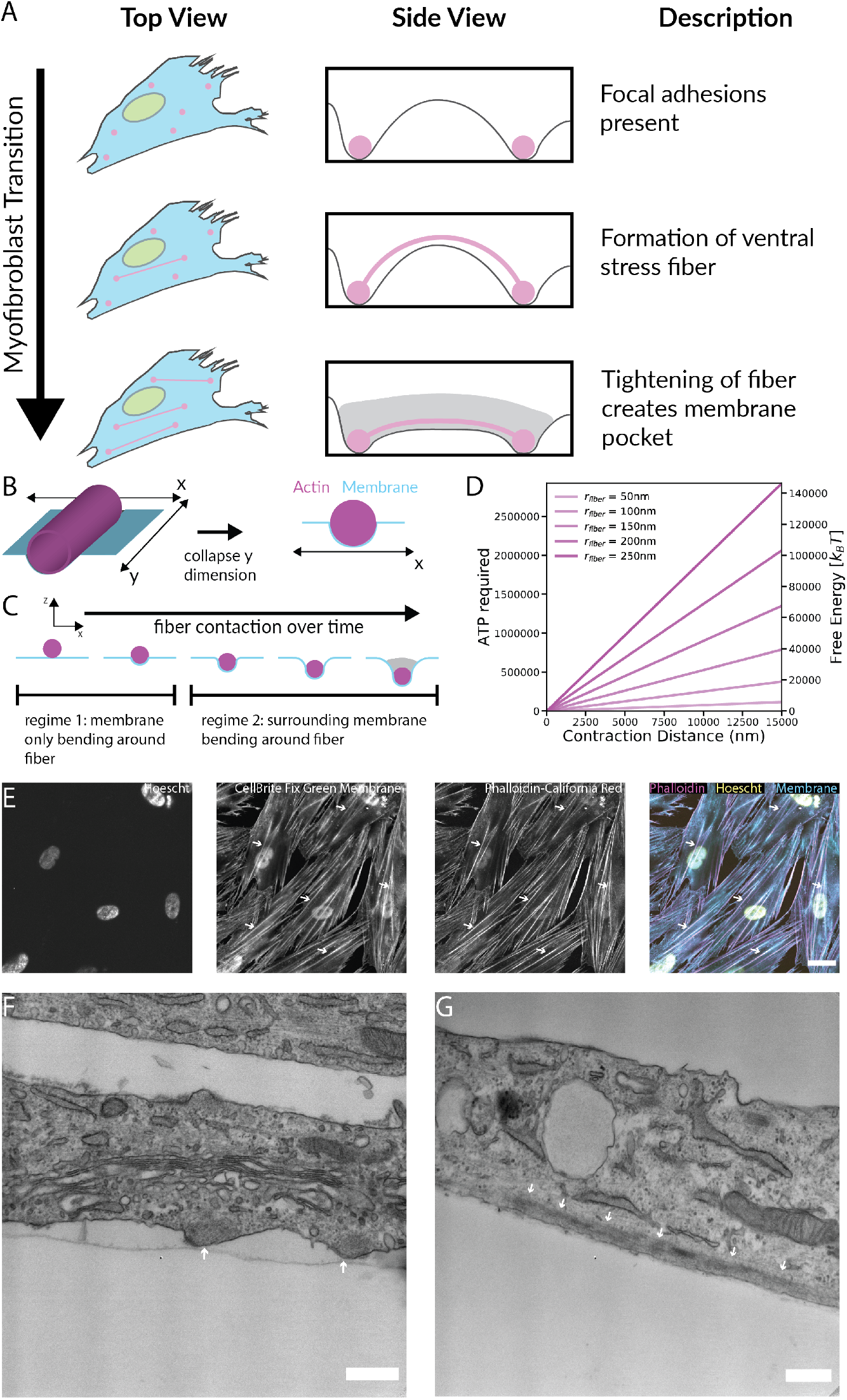
Actin Stress Fibers Induce Membrane Pocket Formation Which can be Labeled with Cytosolic Fluorophores. **(A)** Schematic of a proposed mechanism for the development of the fluorescent structures observed in Figure 1. As fibroblasts transition into myofibrobolasts, ventral actin stress fibers (magenta rods) originating from focal adhesions (magenta circles) deform the plasma membrane, creating cytosolic pockets (grey) for the fluorescent dye or proteins to flow into, leading to the observed fluorescent structures. **(B)** Model conceptualization of stress fiber-induced membrane deformation. The ventral stress fiber is modeled as a cylinder deforming a planar membrane. As the y dimension is uniform, the model is collapsed to one dimension. **(C)** Membrane deformation model used for membrane energy calculation. A stress fiber was modeled as lowering into a membrane, causing the membrane to curve. This fell into two regimes: one where the membrane is only deforming around the fiber and one where parts of the membrane beyond the fiber are deforming. The energy requirement for bending the membrane were calculated across both regimes. **(D)** Calculation results from stress fiber contraction calculation. In our proposed model, the contraction of the ventral stress fibers by myosin II motors drives the formation of the membrane pockets. Here, we consider the energy required for that contraction over a range of observed fiber radii (50-250 nm) and contraction distances (0-15000 nm). The calculated ATP (left y-axis) and *k_B_T* equivalents (right y-axis) indicate the proposed model is reasonable given the timeframe of membrane pocket formation. **(E)** Fluorescent structures can be observed in the Plasma membrane after staining with Green-CellBright Fixable membrane dye (examples marked with white arrows). Like the cytosolic dyes in Figure 2, these fluorescent membrane structures colocalize with ventral actin stress fibers, supporting the hypothesis that ventral actin stress fibers play a role in the formation of the observed fluorescent structure. Scale Bar = 25 *μ*m. **(F)** and **(G)** TEM images of ventral actin stress fibers where **(F)** membrane deformation and **(G)** the thick ventral stress fibers can be directly observed (white arrows). The membrane pockets are distinct from focal adhesions, which appear as a dark plaque near the membrane and exhibit sharper curvature on the edges (Figure S2 and (27–29)). Scale Bar = 500 nm. Note: The cellBrite dye also brightly stains the nuclear membrane, causing some nuclear bleedthrough from the green (cellBrite) channel into the red (phalloidin) channel. Figure 4C replicates this phalloidin/membrane staining with alternate dyes and minimal bleedthrough. Each experiment was conducted in parallel in three separate wells, and a representative image from one well is shown.

To test this hypothesis, we estimated if the forces and energy required to create these ridges are both possible and reasonable within the constraints of the cellular energy budget. To estimate the energy cost of membrane deformation by the ventral stress fiber, we started with calculating the free energy, Gbend, required to bend a membrane:

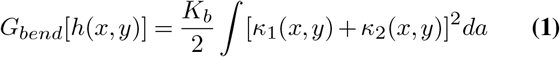

where h(x,y) is the height of the membrane at position (x,y) relative to some reference height, Kb is the membrane bending rigidity (typically on the order of 10-20 *k_B_T*), *κ*_1_ is the curvature of the membrane in the x dimension at position (x,y), *κ*_2_ is the curvature of the membrane in the y dimension at position (x,y), and *da* is differential area. To simplify our calculations, we assumed our fibers to be a straight cylinder indenting a planar membrane (Figure 3B). This collapses the curvature consideration to one dimension, as all of the curvature along the y-axis will be the same at any given x (Figure 3C). This simplified our free energy equation calculation to Equation 2:

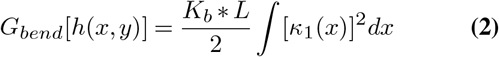

where L is the length of the fiber cylinder. Curvature of a 1D line is calculated using Equation 3:

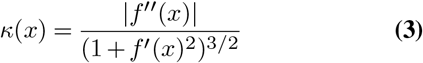

where *f*(*x*) is the function describing the change in height of a line. We built some simple fitting equations to model this profile of the membrane around the fiber (see Methods). The range of free energy requirements fell into two regimes: when the center point of the fiber was modeled above the plane of the membrane and when the center point of the fiber was below the plane of the membrane. In the first regime, the free energy requirements were calculated using the assumption that the membrane bent directly around the fiber (Figure 3C, first two panels). The calculated energies fell in the range of 0-500 *k_B_T*. In the second regime, along with bending around the fiber, parts of the membrane extending out past the fiber diameter were also simulated as bending (Figure 3C, last three panels). While the exact energy values depend on how these bending equations describe the membrane bending, we found the estimates generally fell in the range of a few thousand *k_B_T*. As a comparison point, the free energy of vesicle formation is about 500 *k_B_T*, so the first regime of this membrane bending phenomenon is estimated to be in the same order of magnitude, while the second regime, the one that predicts the pocket where fluorescent molecules would pool (Figure 3C, final panel), falls no more than one order of magnitude above this known phenomenon. After examining the overall energy requirements, we then examined if the actual energy budget required to induce this phenomenon was reasonable given the time frame and estimated energy requirement.

To see if the energy of the membrane bending hypothesis fell within a reasonable energy budget of a fibroblast cell, we estimated the number of ATP molecules required to contract the stress fibers to cause sufficient bending in the membrane. While we recognize that the addition of actin fibers to a ventral stress fiber is a dynamic process, we assumed a static bundle of fibers for the purposes of this calculation. We started by assuming a range of possible radii for the stress fibers. We then calculated the number of individual actin fibers in a given cross-section based on an actin fiber radius of 8 nm. This allowed us to calculate the number of individual fibers of actin that needed to contract using Equation 4:

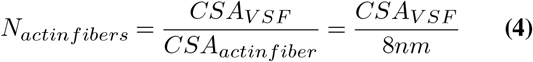

Actin polymers are contracted by myosin motors, whose step size has been estimated at 5 nm (62). It has also been measured that a myosin motor requires 1 ATP/step (63). Using these estimates, we explored a range of ATP requirements for a variety of fiber radii and contraction lengths. Given the assumptions and estimates made, we calculated a linear relationship between energy requirement and how much length the fiber contracts, as shown by Equation 5:

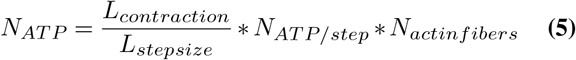

We calculated this relationship over a range of potential radius values for the fibers (Figure 3D). Even for the largest estimate of fiber radius (250 nm), we calculated the energy requirement to be on the order of millions of ATP molecules.

To understand if these values were reasonable, we compared the ATP requirement we calculated with an estimate of the total ATP budget for a fibroblast cell. One calculation estimated fibroblast ATP production at 1 billion ATP/sec/cell (64), putting a multi-day formation of these fibers comfortably within the energy budget of the cell, ensuring that energy constraints were not a reason to rule out our hypothesis that the contraction of ventral stress fibers causes the formation of these structures.

Next, to test our membrane contour hypothesis experimentally, we stained the myofibroblast membranes with Green CellBrite Fix membrane dye (Figure 3E) and the actin stress fibers with phalloidin-California Red. If the cytosolic fluorescent structures were caused by actin stress fibers bending the membrane and forming a cytosolic pocket, we would expect to see similar fluorescent structures in the stained membrane that colocalize with actin stress fibers. Visualizing these stained cells using spinning disk confocal microscopy, we clearly saw fluorescent structures in the cell membrane that colocalize in the xy plane with actin stress fibers, suggesting that the fluorescent structures involve both the membrane as well as the cytosol, supporting our hypothesis. We confirmed these observations by directly visualizing the actin stress fibers and plasma membrane using TEM, with cells sectioned both parallel and perpendicular to the major axis of the cell (Figure 3F&G). In the cross-section perpendicular to major axis of the cell (Figure 3F), curved membrane pockets approximately 1 *μm* in width and 500 nm in height can be seen protruding beneath the main cell body (white arrows). Inside this pocket, there is a 500 nm diameter dark fibril haze, which looks similar to the ventral stress fibers taken in the images parallel to the major axis of the cell (Figure 3G, fiber marked by white arrows) and is consistent with other TEM images of actin stress fibers (5, 65, 66). Importantly, these membrane pockets do not look similar to TEM images of focal adhesions, lacking the distinct dark plaque on the cell membrane characteristic of focal adhesions seen in both our images (Figure S2), and in the literature (27–29). Taken together, our imaging data demonstrates the existence of a cytosolic pocket in the membrane induced by actin stress fibers.

After observing that stress fibers induce membrane contouring, we investigated the structural linkage between ventral stress fibers and the plasma membrane. Paxillin staining (Figure 2B&D) revealed that the ventral actin stress fibers are coupled to the plasma membrane via adapter proteins at focal adhesions. However it also possible for the actin cytoskeleton to bind the membrane at other points via other actin binding proteins, such as Ezrin, Radexin, and Moesin (ERM). In filopodia and membrane ruffles, ERM proteins bind the protruding actin to the membrane (24–26). In epithelial cells undergoing Epithelial to Mesenchymal transition induced by TGF-*β*1 treatment, ERM proteins colocalize with nascent actin stress fibers (67). To see if ERM protein coupling is present in the observed fluorescent ridges, we stained for ERM proteins in our system, in addition to the cytosol with Cell Explorer green and actin stress fibers using California red (Figure 4B). In a separate experiment, we additionally stained for ERM proteins with an Alexa Fluor 514 secondary, the membrane with far red CellBrite Fix membrane dye,the cytosol using Green Cell Explorer cytosolic dye and actin stress fibers using phalloidin-Alexa Fluor 405 (Figure 4 C). In both experiments we observed colocalization between ERM proteins, actin stress fibers, membrane pockets and the cytosolic pocket, suggesting that ERM proteins could be coupling the membrane to the actin stress fibers. Inhibiting ezrin-actin interactions using the ezrin inhibitor NSC66839 (68), caused the membrane pockets to disappear after approximately 4 hours, supporting our hypothesis that active ezrin-actin binding is necessary for membrane contouring (Figure 4D&E). Interestingly, after ezrin inhibition, the actin stress fibers were still visible, but exhibited a curved morphology (Figure 4E), suggesting that the membrane interactions might play a role in maintaining stress fiber persistence length, which could be an interesting topic to explore in future work.

**Fig. 4.**
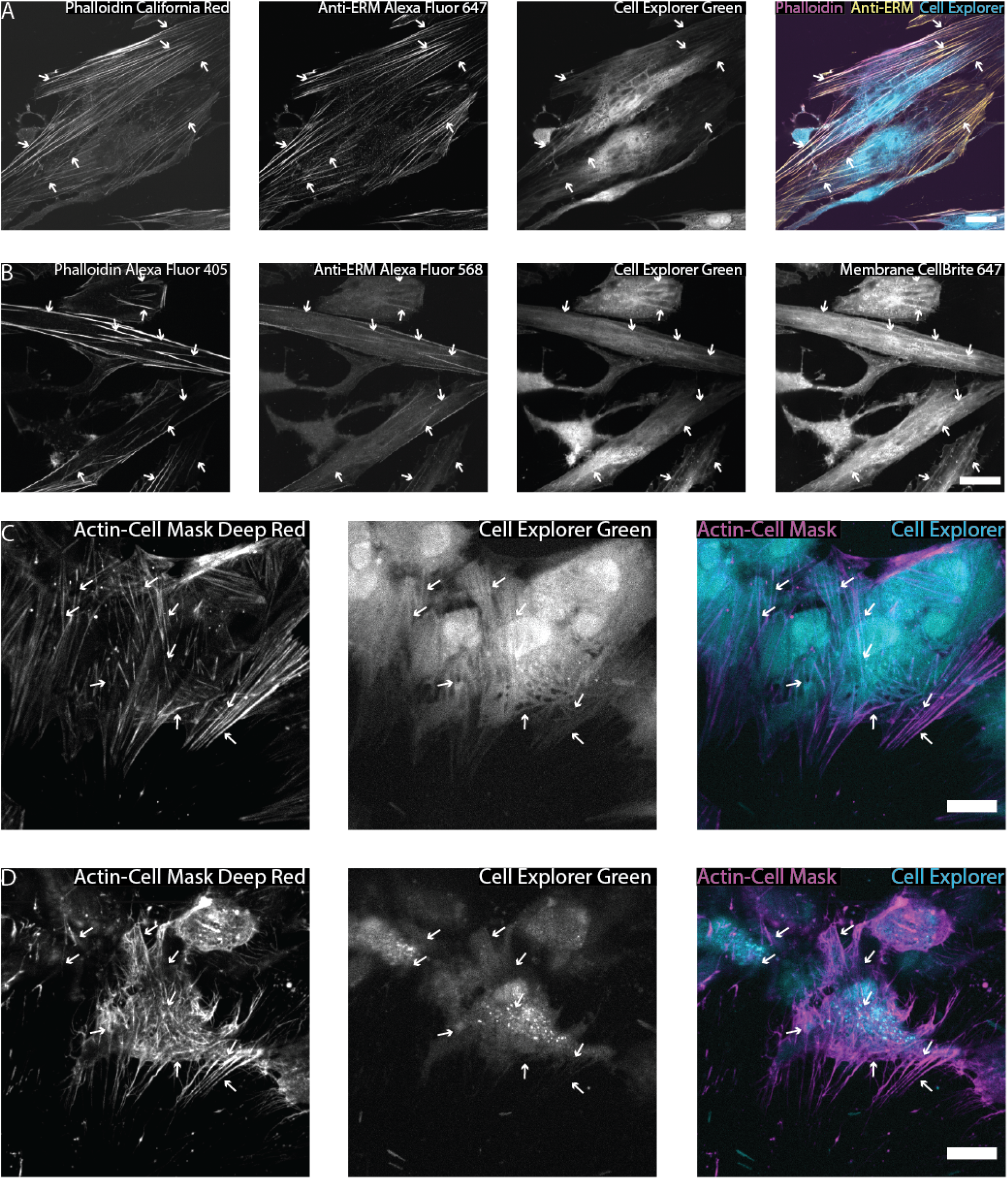
Direct Coupling of Stress Fibers to the Plasma Membrane is Necessary for Membrane Deformation. **(A)** Immunofluorescence staining for ERM proteins reveals that ERM proteins colocalize with stress fibers as well as **(B)** cytosolic and cell membrane pockets. **(C)** Before and **(D)** After images of cells treated with the Ezrin inhibitor NSC66839. Treatment abrogates membrane pockets, but does not dissolve actin fibers, suggesting that physical coupling of the stress fiber to the membrane is necessary for membrane contouring. Scale Bar = 25 *μ*m. Each experiment was conducted in parallel in three separate wells, and a representative image from one well is shown.

Our evidence suggests the membrane is directly coupled to stress fibers along the length of the fiber, so we next investigated the necessity of stress fiber structural integrity for membrane contouring by dissolving the stress fibers. We transfected myofibroblasts with an mCherry-paxillin fusion protein and loaded the cell with Cell Explorer dye and Cell Mask Far Red live cell actin stain to visualize the focal adhesions, cytosolic pocket, and the actin in living cells (5A). We then treated the cells with Cytochalasin-D to disrupt the actin cytoskeleton network (5B). Cytochalasin-D dissolved ventral actin stress fibers along with their associated cytosolic pocket after 15 minutes of drug treatment demonstrating that the physical membrane-stress fiber interaction is necessary for continued membrane contouring. In contrast, focal adhesions (marked by paxillin), which are not dissolved by Cytochalasin-D, and their corresponding cytosolic pocket remained intact, which is consistent with our hypothesis.

### Actin-Induced Membrane Contouring is not Dependent on Cellular Contractility

Myofibroblasts have increased cellular contractility due to their expression of *α-* SMA (51) and the activation of RhoA GTPase (69–72). Cellular contractility derives from contraction of actin stress fibers by the molecular motor myosin II. In particular, the small GTPase RhoA and its downstream Rho associated protein kinase (ROCK) play a key role in cellular contractility by modulating myosin II contraction via myosin light chain phosphotase (MLCP) inhibition (73–76). In our experiments, actin stress fiber deformation of the plasma membrane produces cytosolic pockets, so we investigated the role of actin stress fiber contractility in pocket stability. We stained the cytosol of myofibroblasts using Cell Explorer Green and actin stress fibers using Cell Mask Deep Red actin stain (Figure 5C). We then treated HDFs with 20 uM ROCK inhibitor y-27632 to block RhoA-induced MLCP inhibition and decrease cellular contractility. Y-27632 treatment over 12 hours resulted in actin fiber re-arrangement (Figure 5C&D), but did not remove the associated cytosolic pockets, which moved with the fiber. This suggests that once cytosolic pockets are formed, active stress fiber contraction is not needed to maintain membrane contouring. To further investigate the role of actin contractility in cytosolic pocket stability, we used the drug blebbistatin to directly inhibit myosin II-actin binding. This inhibition of myosin II-actin binding resulted in the disappearance of the cytosolic pocket (Figure 5E&F). This suggests that myosin-mediated actin interactions are necessary for actin fibers to contour the membrane. Taken together, these experiments that disrupt actin structures suggest that once membrane pockets are formed, only structural-stress fiber integrity (such as myosin II-actin binding), and not stress fiber contraction (mediated by ROCK induction of myosin II contraction), is necessary for membrane contouring. However, the role of contractility in membrane pocket formation is still unknown and a promising area for future study.

**Fig. 5.**
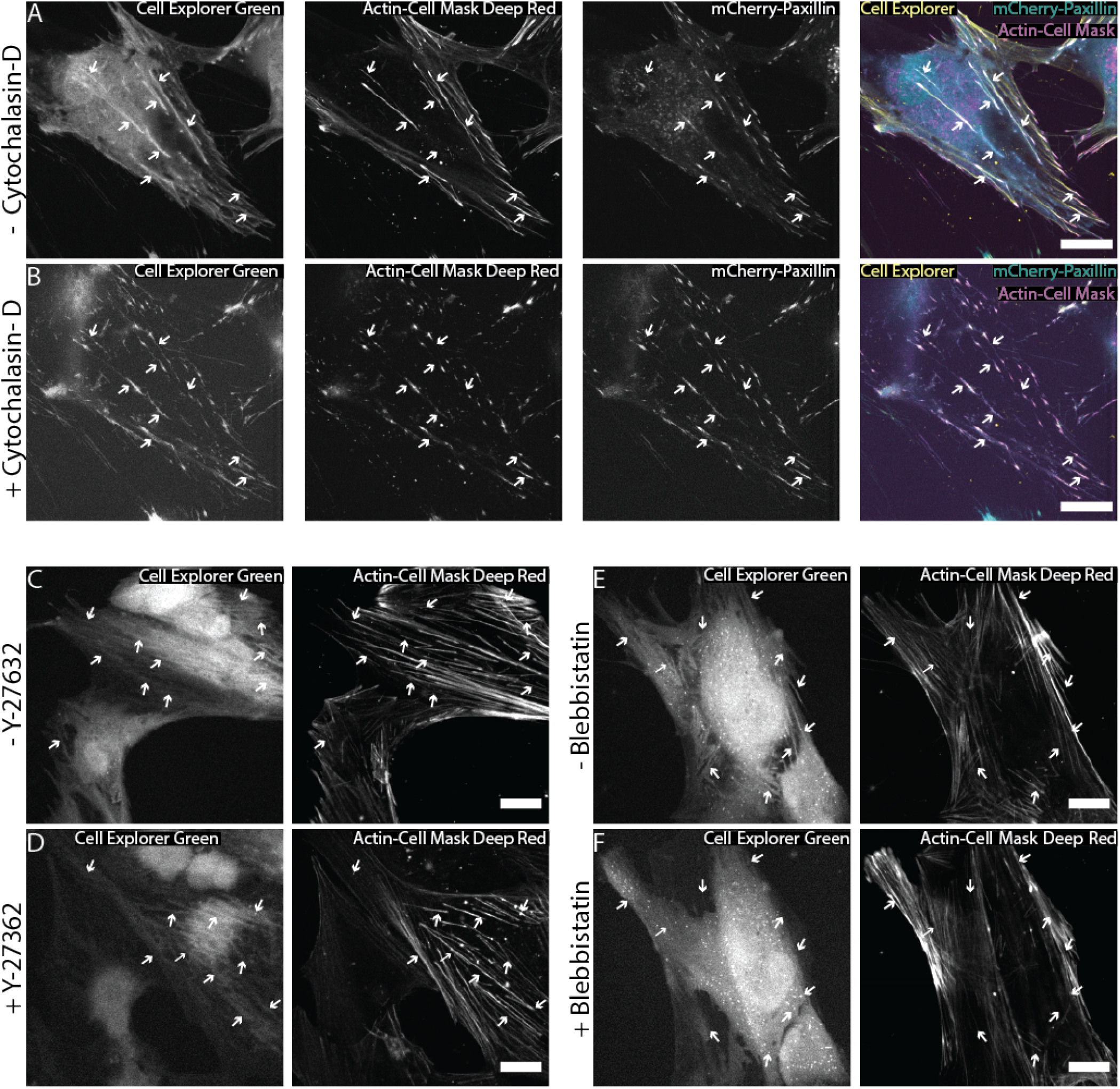
The Structural Integrity, but not Active Contraction of Actin Stress Fibers is Required for Membrane Deformation. **(A)** Before and **(B)** after images of cells treated with Cytochalasin-D. Cytochalasin-D treatment dissolved membrane pockets associated with stress fibers, but not with focal adhesions, indicating that the physical structure of stress fibers is necessary for membrane deformation. **(C)** Before and **(D)** after images of cells treated with Y-27632. Y-27632 treatment had no effect on observed membrane pockets, suggesting that once the membrane has been deformed, stress fiber contraction is not necessary to maintain membrane deformation. **(E)** Before and **(F)** after images of cells treated with Blebbistatin. Blebbistatin treatment also removed observed membrane pockets, suggesting that once the membrane has been deformed, myosin-actin binding is necessary to maintain the membrane pocket. Scale Bar = 25 *μ*m. Each experiment was conducted in parallel in three separate wells, and a representative image from one well is shown.

### Contextualizing Actin Stress Fiber-Plasma Membrane Interactions

In this work, we have described a novel form of actin-induced membrane contouring, where the actin stress fiber applies force perpendicular to the major axis of the fiber. This is in contrast to many actin-membrane interactions, such as those found in filopodia or lamellipodia, where force is applied parallel to the major axis the fiber. In some ways, the new phenomenon is the topological inverse of cytokinesis. During cytokinesis, actin fibers form a ring, with the membrane on the outside of the ring. As the fiber ring contracts, it applies a centripetal force on the membrane, contracting the membrane inward, and eventually splitting the cell in half (Figure 6, left). In the ventral actin case described in this paper, the ventral actin stress fibers can be seen as an arc, with the cell substrate serving as a chord bisecting a larger ring. Here, the membrane is on the inside of the fiber. The fibers then contract and apply the same centripetal force to the membrane, but because the membrane is on the inside of the fiber, it is stretched, rather than contracted, forming the observed membrane pockets (Figure 6, right). There are a few key differences between these two processes. Namely, the actin ring found in cytokinesis is a specialized cytoskeletal structure found in dividing cells (41, 77–80), whereas the ventral actin stress fiber membrane bending occurs with actin stress fibers that primarily exist outside of mitosis (5, 81–83). In addition, cytokinesis requires an active contractile force in the actin ring, either through myosin II contraction (77, 80, 84), actin treadmilling (85, 86), or a combination of both (87). In contrast, our experiments with Y-27632 suggest that once ventral stress fiber associated membrane pockets are observed, active cellular contractility is not needed to maintain membrane bending (Figure 5C-E). While the actin fibers apply force to a membrane in a similar but inverted way to cytokinesis, the phenomena differ in that the ventral actin stress fiber-induced membrane pockets persist without active contraction. The comparison between these two phenomena highlight both the structural and mechanical role actin fibers play in the cell.

**Fig. 6.**
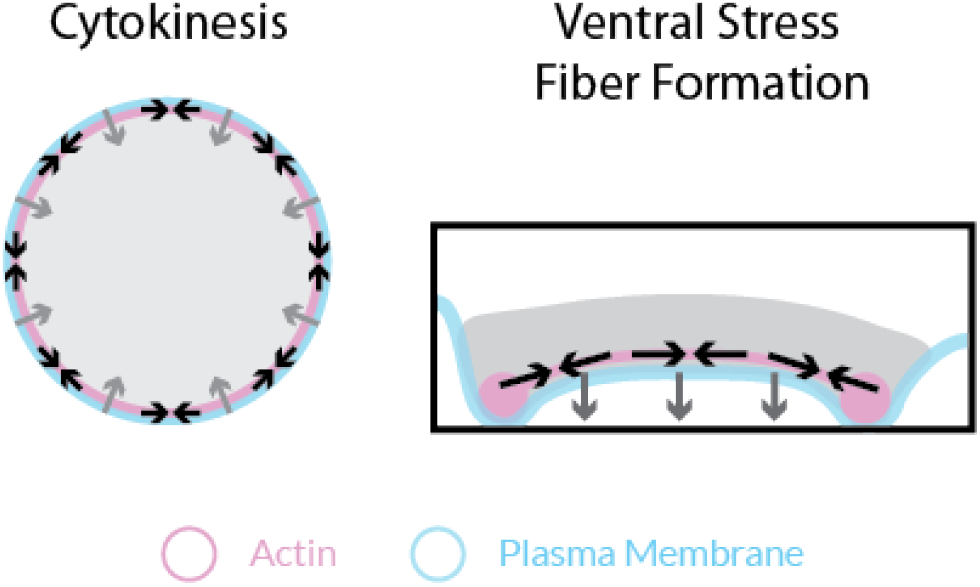
Contextualizing Actin Stress Fiber-Plasma Membrane Interactions. Schematics of cytokinesis **(left)** and lateral membrane bending **(right)**, which demonstrate inverse topologies. In cytokinesis, an actin fiber ring (magenta) contract (black arrows) and apply a centripetal force (gray arrows) to the membrane (cyan) which is on the outside of the fiber, resulting in inward contraction of the membrane, and ultimately membrane cleavage. In the novel ventral stress fiber-induced membrane bending, the ventral stress fibers can be viewed as an arc on a circle. In this case the membrane is on the inside of the fiber, and when these fibers contract (black arrows), and apply the centripetal force to the membrane (gray arrow) expanding the membrane and forming ridges.

The mechanical environment of the cell likely plays a role in stress fiber formation, and thus likely in the formation of the pockets we identified here. There are a few critical differences between 2D cell culture and a 3D tissue environment *in vivo,* namely that in 2D there is artificial cell polarization and often stiffer substrates. By growing myofibroblasts (which are specialized to contract wounds closed (88, 89)) on a very stiff (glass) 2D substrate, we are observing artificially polarized actin-membrane interactions near the extreme end of fibroblast contractility. In 2D culture, the cell can only adhere to a substrate on its ventral side, creating an artificial polarization where focal adhesions only exist on one side of the cell (23, 90), whereas in 3D focal adhesions exist on all sides of the cell (91–94). The forced polarization of focal adhesions in 2D may artificially increase the size and strength of stress fibers compared to stress fibers in 3D (95). In addition, these cells are being grown on glass (Young’s modulus on the order of 70 GPa (96)), which is around 1000 times stiffer than native dermal tissue (Young’s modulus on the order of 70 MPa (97, 98)). Therefore, it is possible that the environmental conditions in 2D that generate the high contractile forces necessary to laterally deform the membrane many not exist in a native 3D environment. However, stress fibers do exist in 3D in *vitro* fibroblast culture (99–101) and in 3D in *vitro* osteoblast culture (102, 103) as well as in 3D in *vivo* tissues including endothelial cell tissues (104, 105) and myofibroblasts in wounded dermal tissues (106–109). The existence of stress fibers in these *in vivo* contexts suggests that, under certain conditions, ventral actin stress fibers could deform the plasma membrane, warranting further investigation into this phenomenon.

## ACKNOWLEDGEMENTS

The authors thank Dr. Emily Mace, Dr. John Ngo and Dr. Emily Hager fortheir critical reviewof our manuscript and helpful comments. This work was supported by the National Institutes of Health (NIGMS) grant R35 GM133616 to A.E.S. M.S.A. was supported by a Multicellular Design Program fellowship from the Rajen Kilachand Fund for Integrated Life Sciences and Engineering. Electron microscopy imaging, consultation, and services were performed in the Harvard Medical School Electron Microscopy Facility.

## Materials and Methods

### Plasmid Construction

pLenti CMV Puro Dest ERK-KTR was digested with *BsrgI* to remove the ERK-KTR gene. The appropriate primers for each gene (see primer table) were used to generate PCR fragments of the gene and add 20-25 base homology arms to each end of the PCR fragment. The digested pLenti CMV Puro Dest vector and PCR fragment were assembled using NEB HiFi Assembly mix, and the mixture was transformed into NEBstable *E. coli* and grown at 30°C overnight. Plasmids were sequence verified by Sanger Sequencing provided by Quintara Biosceince using the inhouse CMV Forward (BP0002) and WPRE (BP0156) reverse primers.

### Viral Production

HEK 293FT cells (passages 3-15) were plated at 90% confluency in a T-25 flask in HEK cell media (DMEM, 10% FBS, 1X Glutamax, 1x NEAA). After 24 hours, the flask was transfected with the pLenti plasmid and the packaging VSV-G and PSPAX-2 plasmids using Lipofec-tamine 3000, according to the manufacturer’s protocol. After 12 hours, the media in the dish was discarded and replaced. Media from the flask containing viral particles was collected 24 hours later, replaced and collected again 24 hours later. The viral media was spun at 300 x g for 10 minutes to pellet any cells, and the supernatant was then passed through a .45 *μ*m syringe filter. The resulting media was aliquoted in 500 *μ*L tubes and stored at −80°C.

### NHDF Cell Culture

Neonatal Human Dermal Fibroblasts (passages 1-8) were cultured in fibroblast media (FGM media supplemented with an FGM-2 OneShot kit) in an incubator at 37°C and 5% CO_2_. Cells were passaged at 80-90% confluency, and media was changed every 48 hours. To generate stably expressing pools of cells, fibroblasts were lifted from a flask by incubating the cells in .05% trypsin and then pelleted by centrifugation at 300 x g for 5 minutes. The cells were then resuspended in fresh fibroblast media and seeded in a 24 well plate at a concentration of 20,000 cells/well. 500 *μL* of viral media and 500 *μL* of fibroblast media were then added to the well, and the dish was incubated at 37 °C for 48 hours. The media in the well was then changed to fibroblast media with 1 *μ*g/mL puromycin to select for positively transduced cells. After 48 hours, the cells were transferred to either a 6 well dish for continued passaging, or a new 24 well plate for experimentation.

### Stress Fiber Induction

Fibroblasts were seeded at 20,000 cells/well in a glass bottomed 24 well dish. After 24 hours, stress fibers were induced by changing the cell media to a serum-free induction media (DMEM, 2% B-27, 10 ng/*μ*L TGĘ*β*-1) which was refreshed every 48 hours. Cells were used after 96 hours of induction.

### Confocal Microscopy

After 96 hours of induction, the cells were stained with various live cell stains according to the manufacturer’s instructions, which generally involved diluting a stock solution 1000x (10,000x for Cell Mask Actin stain) in DMEM, and incubating the cells for 15 - 30 minutes at 37°C and 5% CO_2_. After staining, the cell media was changed to imaging media (Fluorobrite DMEM, 1% Glutamax, 1% OxyFluor). Cells were imaged on a Ti-2E Eclipse (Nikon Instruments) with a Dragonfly Spinning Disk confocal system (Oxford Instruments) in a 37°C and 5% CO_2_ stage top incubator (OKO labs). Images were acquired on an iXon 888 Life EM-CCD camera (Oxford Instruments). Fluorescent dyes were imaged through a 405/488/561/647 dichroic mirror using the following excitation laser/emission filter combinations: Ex.405 nm-Em.445/50, Ex. 488 nm-Em. 515/30, Ex. 561 nm-Em. 590/60, Ex. 647-Em. 698/60. All staining and drug treatments were repeated in 3 separate wells, and a representative image was selected from each treatment for display in a figure.

### Inhibitor Treatments

Cells were treated with the actin inhibitor Cytochalasin-D (5 *μ*M, Tocris), the Ezrin Inhibitor NSC668394 (50 *μ*M, Calbiochem), Y27632 (25 *μ*M, Hello Bio), and s-nitro-Blebbistatin 25 *μ*M, Cayman Chemical). For Cytochalasin-D treatment, an 18 slice z-stack (140 nm step size) was acquired every minute for 5 minutes using a Plan Apochromatic 100x silicone oil immersion objective (Nikon, NA = 1.35) as described above. After 5 minutes, a solution of Cytochalasin-D dissolved in imaging media was injected into the well and the cells were imaged for another 15 minutes post treatment. For NSC668394, Y27632, and S-nitro-Blebbistatin, cells were treated immediately before imaging and an 18 slice z-stack (192.5 nm step size) was acquired every hour for 17 hours using a Plan Apochromatic 40x air objective (Nikon, NA = 0.95) and a 2x zoom lens (80x total magnification). Each drug treatment experiment was repeated in three separate wells and multiple fields of view were collected per well.

### Immunofluorescence

After 96 hours of induction, cells were fixed in 4% Paraformaldehyde in PBS and permeabilized using 0.1% Triton-x. Non-specific interactions were blocked using 10% normal goat serum in PBS. The cells were then incubated overnight with the primary antibody (anti-paxillin 1:50 and anti-ERM 1:100), washed in PBS, and then followed by incubation with the appropriate Alexa Fluor 647 secondary antibody (1:200) in 10% normal goat serum for 1 hour at room temperature. In experiments where phalloidin or membrane stain was used, it was added after the secondary at 1:1000 in PBS and incubated for 30 minutes at room temperature. Cells were washed 3X in PBS and then imaged using the same parameters as previously described using a Plan Apochromatic 100x silicone oil immersion objective (Nikon), but at room temperature with no CO_2_.

### Transmission Electron Microscopy

After 96 hours of induction, cells were fixed in 2.5% paraformaldehyde/glutaraldehyde in sodium cacodylate buffer for 1 hour at room temperature. Cells were then washed with sodium cacodylate buffer 3x and stored at 4 °C until imaged. Electron microscopy imaging, consultation, and services were performed in the HMS Electron Microscopy Facility, on a TecnaiG2Spirit BioTwin microscope with a 2k AMT camera.

### Membrane Bending Energy Calculations

All model calculations were carried out using a Python script in Spyder (version 4.1.5) using the NumPy (110), Matplotlib (111), and SciPy (112) packages. To calculate membrane bending, the profile of the membrane wrapping around the fiber was defined as follows: until the midpoint of the fiber is level with the membrane, the membrane wraps directly around the fiber using a square root function. As the midpoint of the fiber dips below the level of the membrane, parts of the membrane to either side begin to bend. This is modeled by fitting hyperbolic curves that start at the y value at the midpoint of the fiber and have a length half the distance from the fiber midpoint to the resting level of the membrane. After the profile of the membrane has been defined, the curvature at each point was calculated using the derivative function from the SciPy package. The bending energy equation is applied for each point of curvature and summed over the whole stretch of membrane modeled to get the total bending energy. Code available online at https://github.com/sgrolab/ventralsfpaper.

## Cells

Cells.
| Cell Name | Manufacturer | Lot |
| --- | --- | --- |
| Neonatal Human Dermal Fibroblasts | Lonza (CC-2509) | 0000490825, 0000490827 |
| HEK293FT | Thermo Fisher (R70007) |  |

## Plasmids

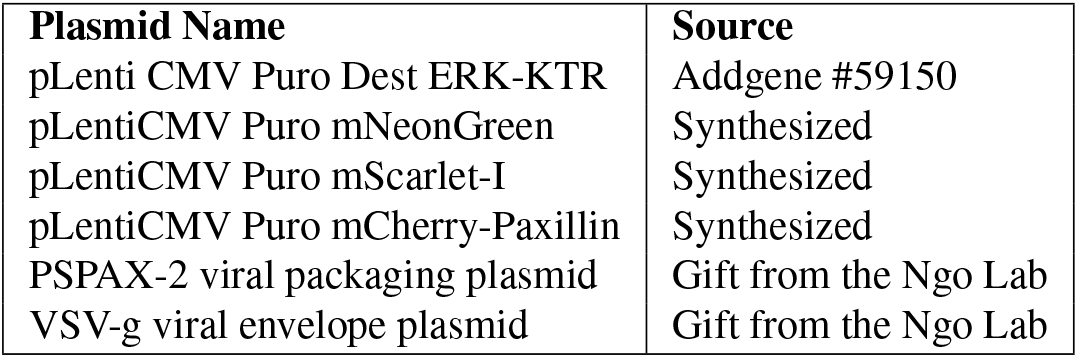

Plasmids.
| Plasmid Name | Source |
| --- | --- |
| pLenti CMV Puro Dest ERK-KTR | Addgene #59150 |
| pLentiCMV Puro mNeonGreen | Synthesized |
| pLentiCMV Puro mScarlet-I | Synthesized |
| pLentiCMV Puro mCherry-Paxillin | Synthesized |
| PSPAX-2 viral packaging plasmid | Gift from the Ngo Lab |
| VSV-g viral envelope plasmid | Gift from the Ngo Lab |

## Reagents

### A. Cell Culture

Reagents. A. Cell Culture.
| Name | Manufacturer | Lot |
| --- | --- | --- |
| FBM media | Lonza (CC-3131) | 0000849191, 0000914535 |
| FGM-2 OneShot | Lonza (CC-4126) | 0000851116, 0000885722, 0000923831 |
| DMEM | Corning (MT10013CV) | 10420009 |
| Opti-MEM | Gibco (31985-062) | 1897019 |
| FluoroBrite DMEM | Gibco (a18967-01) | 2120559 |
| Fetal Bovine Serum | Gibco (16000044) | 2103017RP |
| Pen-Strep | Gibco (16140-122) | 2145104 |
| B-27 | Gibco (17504044) | 2121033, 2193555 |
| Glutamax | Gibco (35050-061) | 2164667 |
| Lipofectamine 3000 | Thermo Fisher (L3000) | 2177271 |
| .05% Trypsin | Gibco (253000-054) | 2120736 |
| OxyFluor | Sigma-Aldrich (SAE0059) | SLCC2576 |
| 24 Well Glass Bottom Dishes | Porvair Sciences (324042) | 037505 |
| 24 Well Plastic Dishes | Denville Scientific (T1024) | 2018003 |
| T-75 culture flasks | Fisher Scientific (156499) | 161855 |
| T-25 culture flasks | Denville Scientific (T1205) | 2019001 |

### B. Drugs

B. Drugs.
| Name | Manufacturer | Lot |
| --- | --- | --- |
| Recombinant TGFβ-1 | R&D Systems (7754-BH) | DCPU0819111 |
| Cytochalasin-D | Tocris (1233) | 7A/207303 |
| Y-27632 | Hello Bio (HB2297) | E0807-1-4 |
| NSC668394 | calbiochem (341216) | 3434830 |
| s-Nitro-Blebbistatin | Cayman Chemical Company (13891) |  |

### C. Primers for Cloning

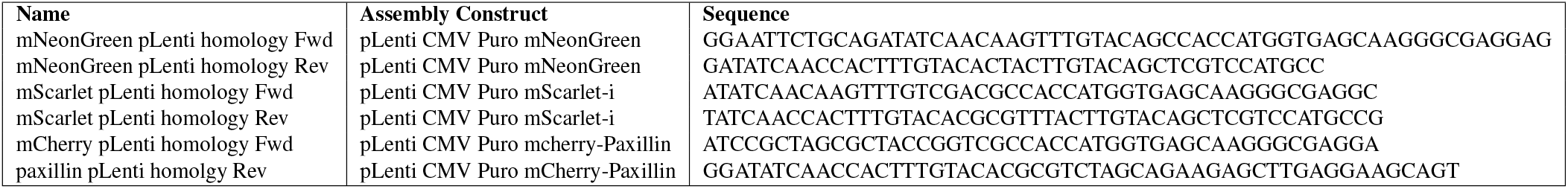

### D. Immunofluorescence

**D. Immunofluorescence.**
| **Name** | **Manufacturer** | **Lot** |
| --- | --- | --- |
| 16% PFA | Thermo Fisher (PI28906) | UG287039 |
| PBS | Gibco (7011-044) | 2193380 |
| Triton-x | Fisher (BP151) | 176408 |
| Normal Goat Serum | Southern Biotech (0060-01) | F3320-SD30B |
| Phalloidin Alexa Fluor 405 | Thermo Fisher (A30104) | (0529804-1) |
| Phalloidin iFluor 488 | Cayman Chemical (20549) | 0593927 |
| Phalloidin California Red | Cayman Chemical (20546) | 0529804 |
| Rabbit anti-paxillin antibody | Cell Signaling Technology (69363S) | 1 |
| Rabbit anti-ERM antibody | Cell Signaling Technology (3142T) | 5 |
| Goat anti-mouse Alexa Fluor 647 conjugate | Thermo Fisher (A21244) | 2161043 |
| Goat anti-rabbit Alexa Fluor 647 conjugate | Thermo Fisher (A21235) | 2134003 |
| Goat anti-rabbit Alexa Fluor 568 conjugate | Abcam (175471) | GR3189790-3 |
| Hoechst 34580 | Sigma Aldrich (63493) | BCCB3802 |
| Cell Explorer dye Green | AAT Bioquest (22621) | 2391532 |
| CellBrite Fix 640 membrane dye | Biotium (30089T) | 17C1115-1104173 |
| CellBrite Fix 488 membrane dye | Biotium (30090A) | 18C0118-1120003 |
| CellMask™ Deep Red Actin Tracking Stain | Thermo Fisher (A57245) | 2212430 |

### E. Plasmid Cloning

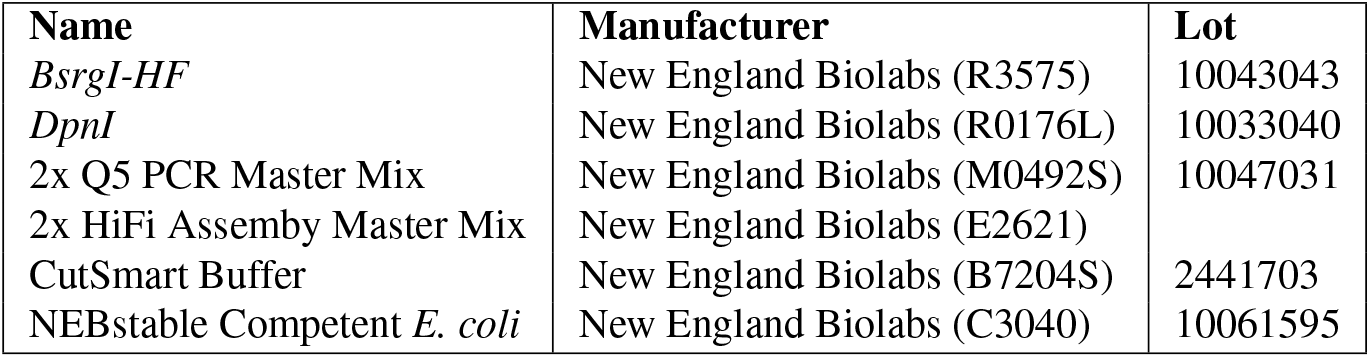

**E. Plasmid Cloning.**
| **Name** | **Manufacturer** | **Lot** |
| --- | --- | --- |
| *BsrgI-HF* | New England Biolabs (R3575) | 10043043 |
| *DpnI* | New England Biolabs (R0176L) | 10033040 |
| 2x Q5 PCR Master Mix | New England Biolabs (M0492S) | 10047031 |
| 2x HiFi Assembly Master Mix | New England Biolabs (E2621) |  |
| CutSmart Buffer | New England Biolabs (B7204S) | 2441703 |
| NEBstable Competent *E. coli* | New England Biolabs (C3040) | 10061595 |

### F. PTEM Prep

**F. TEM Prep.**
| **Name** | **Manufacturer** | **Lot** |
| --- | --- | --- |
| Formaldehyde/Glutaraldehyde 2.5% | Electron Microscopy Sciences (NC0620709) | 200928-04 |
| Sodium Cacodylate Buffer | Electron Microscopy Sciences (11650) | 200811 |
| 12 mm CoverSlips | Fisher Scientific (C3040) |  |

## Bibliography

1. Chun-Min Lo, Hong-Bei Wang, Micah Dembo, and Yu-li Wang. Cell Movement Is Guided by the Rigidity of the Substrate. Biophysical Journal, 79(1):144–152, July 2000. doi: 10.1016/S0006-3495(00)76279-5.

2. Shang-You Tee, Jianping Fu, Christopher S. Chen, and Paul A. Janmey. Cell Shape and Substrate Rigidity Both Regulate Cell Stiffness. Biophysical Journal, 100(5):L25–L27, March 2011. doi: 10.1016/j.bpj.2010.12.3744.

3. Dennis W. Zhou, Ted T. Lee, Shinuo Weng, Jianping Fu, and Andrés J. García. Effects of substrate stiffness and actomyosin contractility on coupling between force transmission and vinculin-paxillin recruitment at single focal adhesions. Molecular Biology of the Cell, 28(14):1901–1911, May 2017. doi: 10.1091/mbc.e17-02-0116.

4. L. P Cramer. Organization and polarity of actin filament networks in cells: implications for the mechanism of myosin-based cell motility. Biochemical Society Symposium, 65: 173–205, 1999.

5. Louise P. Cramer, Margaret Siebert, and Timothy J. Mitchison. Identification of Novel Graded Polarity Actin Filament Bundles in Locomoting Heart Fibroblasts: Implications for the Generation of Motile Force. Journal of Cell Biology, 136(6):1287–1305, March 1997. doi: 10.1083/jcb.136.6.1287.

6. Muhammad H. Zaman, Linda M. Trapani, Alisha L. Sieminski, Drew MacKellar, Haiyan Gong, Roger D. Kamm, Alan Wells, Douglas A. Lauffenburger, and Paul Matsudaira. Migration of tumor cells in 3D matrices is governed by matrix stiffness along with cell-matrix adhesion and proteolysis. Proceedings of the National Academy of Sciences, 103(29): 10889–10894, July 2006. doi: 10.1073/pnas.0604460103.

7. Jianping Fu, Yang-Kao Wang, Michael T. Yang, Ravi A. Desai, Xiang Yu, Zhijun Liu, and Christopher S. Chen. Mechanical regulation of cell function with geometrically modulated elastomeric substrates. Nature Methods, 7(9):733–736, September 2010. doi: 10.1038/nmeth.1487.

8. Ying Wang and Veit Riechmann. The Role of the Actomyosin Cytoskeleton in Coordination of Tissue Growth during Drosophila Oogenesis. Current Biology, 17(15):1349–1355, August 2007. doi: 10.1016/j.cub.2007.06.067.

9. Kandice R. Levental, Hongmei Yu, Laura Kass, Johnathon N. Lakins, Mikala Egeblad, Janine T. Erler, Sheri F. T. Fong, Katalin Csiszar, Amato Giaccia, Wolfgang Weninger, Mitsuo Yamauchi, David L. Gasser, and Valerie M. Weaver. Matrix Crosslinking Forces Tumor Progression by Enhancing Integrin Signaling. Cell, 139(5):891–906, November 2009. doi: 10.1016/j.cell.2009.10.027.

10. Hong-Bei Wang, Micah Dembo, and Yu-Li Wang. Substrate flexibility regulates growth and apoptosis of normal but not transformed cells. American Journal of Physiology-Cell Physiology, 279(5):C1345–C1350, November 2000. doi: 10.1152/ajpcell.2000.279.5.C1345.

11. Adam J. Engler, Shamik Sen, H. Lee Sweeney, and Dennis E. Discher. Matrix Elasticity Directs Stem Cell Lineage Specification. Cell, 126(4):677–689, August 2006. doi: 10.1016/j.cell.2006.06.044.

12. Robert J. Pelham and Yu-li Wang. Cell locomotion and focal adhesions are regulated by substrate flexibility. Proceedings of the National Academy of Sciences, 94(25):13661–13665, December 1997. doi: 10.1073/pnas.94.25.13661.

13. Julien Colombelli, Achim Besser, Holger Kress, Emmanuel G. Reynaud, Philippe Girard, Emmanuel Caussinus, Uta Haselmann, John V. Small, Ulrich S. Schwarz, and Ernst H. K. Stelzer. Mechanosensing in actin stress fibers revealed by a close correlation between force and protein localization. Journal of Cell Science, 122(11):1928–1928, June 2009. doi: 10.1242/jcs.054577.

14. Fabiana Martino, Ana R. Perestrelo, Vladimír Vinarský, Stefania Pagliari, and Giancarlo Forte. Cellular Mechanotransduction: From Tension to Function. Frontiers in Physiology, 9, 2018. doi: 10.3389/fphys.2018.00824.

15. Michele A. Wozniak and Christopher S. Chen. Mechanotransduction in development: a growing role for contractility. Nature Reviews Molecular Cell Biology, 10(1):34–43, January 2009. doi: 10.1038/nrm2592.

16. Christopher S. Chen. Mechanotransduction - a field pulling together? Journal of Cell Science, 121 (20):3285, October 2008. doi: 10.1242/jcs.023507.

17. K. Weber and U. Groeschel-Stewart. Antibody to Myosin: The Specific Visualization of Myosin-Containing Filaments in Nonmuscle Cells. Proceedings of the National Academy of Sciences, 71(11):4561–4564, November 1974. doi: 10.1073/pnas.71.11.4561.

18. A. B. Verkhovsky and G. G. Borisy. Non-sarcomeric mode of myosin II organization in the fibroblast lamellum. Journal of Cell Biology, 123(3):637–652, November 1993. doi: 10.1083/jcb.123.3.637.

19. Todd Thoresen, Martin Lenz, and Margaret L. Gardel. Reconstitution of Contractile Actomyosin Bundles. Biophysical Journal, 100(11):2698–2705, June 2011. doi: 10.1016/j.bpj.2011.04.031.

20. Jonathan D. Humphries, Pengbo Wang, Charles Streuli, Benny Geiger, Martin J. Humphries, and Christoph Ballestrem. Vinculin controls focal adhesion formation by direct interactions with talin and actin. The Journal of Cell Biology, 179(5):1043–1057, December 2007. doi: 10.1083/jcb.200703036.

21. Michio Muguruma, Sueo Matsumura, and Toshiyuki Fukazawa. Direct interactions between talin and actin. Biochemical and Biophysical Research Communications, 171(3): 1217–1223, September 1990. doi: 10.1016/0006-291X(90)90815-5.

22. David A. Calderwood, Yosuke Fujioka, Jose M. de Pereda, Begoña García-Alvarez, Tetsuya Nakamoto, Ben Margolis, C. Jane McGlade, Robert C. Liddington, and Mark H. Ginsberg. Integrin β cytoplasmic domain interactions with phosphotyrosine-binding domains: A structural prototype for diversity in integrin signaling. Proceedings of the National Academy of Sciences, 100(5):2272–2277, March 2003. doi: 10.1073/pnas.262791999.

23. Pirta Hotulainen and Pekka Lappalainen. Stress fibers are generated by two distinct actin assembly mechanisms in motile cells. Journal of Cell Biology, 173(3):383–394, May 2006. doi: 10.1083/jcb.200511093.

24. Monique Arpin, Marianne Algrain, and Daniel Louvard. Membrane-actin microfilament connections: an increasing diversity of players related to band 4.1. Current Opinion in Cell Biology, 6(1):136–141, February 1994. doi: 10.1016/0955-0674(94)90127-9.

25. O. Turunen, T. Wahlström, and A. Vaheri. Ezrin has a COOH-terminal actin-binding site that is conserved in the ezrin protein family. Journal of Cell Biology, 126(6):1445–1453, September 1994. doi: 10.1083/jcb.126.6.1445.

26. Nao Nakamura, Noriko Oshiro, Yuko Fukata, Mutsuki Amano, Masaki Fukata, Shinya Kuroda, Yoshiharu Matsuura, Thomas Leung, Louis Lim, and Kozo Kaibuchi. Phosphorylation of ERM proteins at filopodia induced by Cdc42. Genes to Cells, 5(7):571–581, 2000. doi: 10.1046/j.1365-2443.2000.00348.x.

27. M. Abercrombie, Joan E. M. Heaysman, and Susan M. Pegrum. The locomotion of fibroblasts in culture: IV. Electron microscopy of the leading lamella. Experimental Cell Research, 67(2):359–367, August 1971. doi: 10.1016/0014-4827(71)90420-4.

28. Ohad Medalia and Benjamin Geiger. Frontiers of microscopy-based research into cell-matrix adhesions. Current Opinion in Cell Biology, 22(5):659–668, October 2010. doi: 10.1016/j.ceb.2010.08.006.

29. M. Abercrombie, Joan E. M. Heaysman, and Susan M. Pegrum. The locomotion of fibroblasts in culture I. Movements of the leading edge. Experimental Cell Research, 59(3): 393–398, March 1970. doi: 10.1016/0014-4827(70)90646-4.

30. Tatyana Svitkina. The Actin Cytoskeleton and Actin-Based Motility. Cold Spring Harbor Perspectives in Biology, 10(1):a018267, January 2018. doi: 10.1101/cshperspect.a018267.

31. J. V. Small and J. E. Celis. Filament arrangements in negatively stained cultured cells: the organization of actin. Cytobiologie, 16(2):308–325, February 1978.

32. Albrecht Wegner. Head to tail polymerization of actin. Journal of Molecular Biology, 108 (1):139–150, November 1976. doi: 10.1016/S0022-2836(76)80100-3.

33. J. D. Cortese, B. Schwab, C. Frieden, and E. L. Elson. Actin polymerization induces a shape change in actin-containing vesicles. Proceedings of the National Academy of Sciences, 86(15):5773–5777, August 1989. doi: 10.1073/pnas.86.15.5773.

34. A. Mogilner and G. Oster. Cell motility driven by actin polymerization. Biophysical Journal, 71(6):3030–3045, December 1996. doi: 10.1016/S0006-3495(96)79496-1.

35. Grégory Giannone, Benjamin J. Dubin-Thaler, Olivier Rossier, Yunfei Cai, Oleg Chaga, Guoying Jiang, William Beaver, Hans-Günther Döbereiner, Yoav Freund, Gary Borisy, and Michael P. Sheetz. Lamellipodial Actin Mechanically Links Myosin Activity with Adhesion-Site Formation. Cell, 128(3):561–575, February 2007. doi: 10.1016/j.cell.2006.12.039.

36. Erin M. Craig, David Van Goor, Paul Forscher, and Alex Mogilner. Membrane Tension, Myosin Force, and Actin Turnover Maintain Actin Treadmill in the Nerve Growth Cone. Biophysical Journal, 102(7):1503–1513, April 2012. doi: 10.1016/j.bpj.2012.03.003.

37. Caspar Rüegg, Claudia Veigel, Justin E. Molloy, Stephan Schmitz, John C. Sparrow, and Rainer H. A. Fink. Molecular Motors: Force and Movement Generated by Single Myosin II Molecules. Physiology, 17(5):213–218, October 2002. doi: 10.1152/nips.01389.2002.

38. Miguel Vicente-Manzanares, Xuefei Ma, Robert S. Adelstein, and Alan Rick Horwitz. Nonmuscle myosin II takes centre stage in cell adhesion and migration. Nature Reviews Molecular Cell Biology, 10(11):778–790, November 2009. doi: 10.1038/nrm2786.

39. I. Mabuchi and M. Okuno. The effect of myosin antibody on the division of starfish blastomeres. Journal of Cell Biology, 74(1):251–263, July 1977. doi: 10.1083/jcb.74.1.251.

40. Robert J. Pelham and Fred Chang. Actin dynamics in the contractile ring during cytokinesis in fission yeast. Nature, 419(6902):82–86, September 2002. doi: 10.1038/nature00999.

41. Dhivya Subramanian, Junqi Huang, Mayalagu Sevugan, Robert C. Robinson, Mohan K. Balasubramanian, and Xie Tang. Insight into Actin Organization and Function in Cytokinesis from Analysis of Fission Yeast Mutants. Genetics, 194(2):435–446, June 2013. doi: 10.1534/genetics.113.149716.

42. Kendall Powell. Myosin powers cytokinesis. Journal of Cell Biology, 170(4):515–515, August 2005. doi: 10.1083/jcb1704fta3.

43. R. Dyche Mullins, John A. Heuser, and Thomas D. Pollard. The interaction of Arp2/3 complex with actin: Nucleation, high affinity pointed end capping, and formation of branching networks of filaments. Proceedings of the National Academy of Sciences, 95(11):6181–6186, May 1998. doi: 10.1073/pnas.95.11.6181.

44. R. Dyche Mullins, Walter F. Stafford, and Thomas D. Pollard. Structure, Subunit Topology, and Actin-binding Activity of the Arp2/3 Complex from Acanthamoeba. Journal of Cell Biology, 136(2):331–343, January 1997. doi: 10.1083/jcb.136.2.331.

45. Praveen Suraneni, Boris Rubinstein, Jay R. Unruh, Michael Durnin, Dorit Hanein, and Rong Li. The Arp2/3 complex is required for lamellipodia extension and directional fibroblast cell migration. Journal of Cell Biology, 197(2):239–251, April 2012. doi: 10.1083/jcb.201112113.

46. Kurt J. Amann and Thomas D. Pollard. Direct real-time observation of actin filament branching mediated by Arp2/3 complex using total internal reflection fluorescence microscopy. Proceedings of the National Academy of Sciences, 98(26):15009–15013, December 2001. doi: 10.1073/pnas.211556398.

47. Laura M. Machesky and Robert H. Insall. Scar1 and the related Wiskott-Aldrich syndrome protein, WASP, regulate the actin cytoskeleton through the Arp2/3 complex. Current Biology, 8(25):1347–1356, December 1998. doi: 10.1016/S0960-9822(98)00015-3.

48. Congying Wu, Sreeja B. Asokan, Matthew E. Berginski, Elizabeth M. Haynes, Norman E. Sharpless, Jack D. Griffith, Shawn M. Gomez, and James E. Bear. Arp2/3 Is Critical for Lamellipodia and Response to Extracellular Matrix Cues but Is Dispensable for Chemotaxis. Cell, 148(5):973–987, March 2012. doi: 10.1016/j.cell.2011.12.034.

49. Boris Hinz, Sem H. Phan, Victor J. Thannickal, Andrea Galli, Marie-Luce Bochaton-Piallat, and Giulio Gabbiani. The Myofibroblast: One Function, Multiple Origins. The American Journal of Pathology, 170(6):1807–1816, June 2007. doi: 10.2353/ajpath.2007.070112.

50. A. Moustakas and C. Stournaras. Regulation of actin organisation by TGF-beta in H-ras-transformed fibroblasts. Journal of Cell Science, 112(8):1169, April 1999.

51. A. Desmoulière, A. Geinoz, F. Gabbiani, and G. Gabbiani. Transforming growth factor-beta 1 induces alpha-smooth muscle actin expression in granulation tissue myofibroblasts and in quiescent and growing cultured fibroblasts. Journal of Cell Biology, 122(1):103–111, July 1993. doi: 10.1083/jcb.122.1.103.

52. Boris Hinz, Giuseppe Celetta, James J. Tomasek, Giulio Gabbiani, and Christine Chapon-nier. Alpha-Smooth Muscle Actin Expression Upregulates Fibroblast Contractile Activity. Molecular Biology of the Cell, 12(9):2730–2741, September 2001. doi: 10.1091/mbc.12.9.2730.

53. Daphne S. Bindels, Lindsay Haarbosch, Laura van Weeren, Marten Postma, Katrin E. Wiese, Marieke Mastop, Sylvain Aumonier, Guillaume Gotthard, Antoine Royant, Mark A. Hink, and Theodorus W. J. Gadella. mScarlet: a bright monomeric red fluorescent protein for cellular imaging. Nature Methods, 14(1):53–56, January 2017. doi: 10.1038/nmeth.4074.

54. Dawen Cai, Kimberly B. Cohen, Tuanlian Luo, Jeff W. Lichtman, and Joshua R. Sanes. Improved tools for the Brainbow toolbox. Nature Methods, 10(6):540–547, June 2013. doi: 10.1038/nmeth.2450.

55. Keith Burridge and Christophe Guilluy. Focal adhesions, stress fibers and mechanical tension. Experimental Cell Research, 343(1):14–20, April 2016. doi: 10.1016/j.yexcr.2015.10.029.

56. K. Burridge, L. Molony, and T. Kelly. Adhesion Plaques: Sites of Transmembrane Interaction between the Extracellular Matrix and the Actin Cytoskeleton. Journal of Cell Science, 1987(Supplement 8):211–219, March 1987. doi: 10.1242/jcs.1987.Supplement_8.12.

57. J.Victor Small, K. Rottner, I. Kaverina, and K.I. Anderson. Assembling an actin cytoskeleton for cell attachment and movement. Biochimica et Biophysica Acta (BBA) - Molecular Cell Research, 1404(3):271–281, September 1998. doi: 10.1016/S0167-4889(98)00080-9.

58. Ariel Livne and Benjamin Geiger. The inner workings of stress fibers from contractile machinery to focal adhesions and back. Journal of Cell Science, 129(7):1293–1304, April 2016. doi: 10.1242/jcs.180927.

59. Nathan C. Shaner, Gerard G. Lambert, Andrew Chammas, Yuhui Ni, Paula J. Cranfill, Michelle A. Baird, Brittney R. Sell, John R. Allen, Richard N. Day, Maria Israelsson, Michael W. Davidson, and Jiwu Wang. A bright monomeric green fluorescent protein derived from Branchiostoma lanceolatum. Nature Methods, 10(5):407–409, May 2013. doi: 10.1038/nmeth.2413.

60. Julia Riedl, Alvaro H. Crevenna, Kai Kessenbrock, Jerry Haochen Yu, Dorothee Neukirchen, Michal Bista, Frank Bradke, Dieter Jenne, Tad A. Holak, Zena Werb, Michael Sixt, and Roland Wedlich-Soldner. Lifeact: a versatile marker to visualize F-actin. Nature Methods, 5(7):605–607, July 2008. doi: 10.1038/nmeth.1220.

61. Michael J. Schell, Christophe Erneux, and Robin F. Irvine. Inositol 1,4,5-Trisphosphate 3-Kinase A Associates with F-actin and Dendritic Spines via Its N Terminus. Journal of Biological Chemistry, 276(40):37537–37546, October 2001. doi: 10.1074/jbc.M104101200.

62. Nikolas Hundt, Walter Steffen, Salma Pathan-Chhatbar, Manuel H. Taft, and Dietmar J. Manstein. Load-dependent modulation of non-muscle myosin-2A function by tropomyosin 4.2. ScientificReports, 6(1):20554, February 2016. doi: 10.1038/srep20554.

63. M. Rief, R. S. Rock, A. D. Mehta, M. S. Mooseker, R. E. Cheney, and J. A. Spudich. Myosin-V stepping kinetics: A molecular model for processivity. Proceedings of the National Academy of Sciences, 97(17):9482–9486, August 2000. doi: 10.1073/pnas.97.17.9482.

64. Avi Flamholz, Rob Phillips, and Ron Milo. The quantified cell. Molecular Biology of the Cell, 25(22):3497–3500, November 2014. doi: 10.1091/mbc.e14-09-1347.

65. J. F. Perdue. THE DISTRIBUTION, ULTRASTRUCTURE, AND CHEMISTRY OF MICROFILAMENTS IN CULTURED CHICK EMBRYO FIBROBLASTS. The Journal of Cell Biology, 58(2):265–283, August 1973. doi: 10.1083/jcb.58.2.265.

66. N. Scott McNutt, Lloyd A. Culp, and Paul H. Black. CONTACT-INHIBITED REVERTANT CELL LINES ISOLATED FROM SV40-TRANSFORMED CELLS II. Ultrastructural Study. Journal of Cell Biology, 50(3):691–708, September 1971. doi: 10.1083/jcb.50.3.691.

67. Stephen T. Buckley, Carlos Medina, Michael Kasper, and Carsten Ehrhardt. Interplay between RAGE, CD44, and focal adhesion molecules in epithelial-mesenchymal transition of alveolar epithelial cells. American Journal of Physiology-Lung Cellular and Molecular Physiology, 300(4):L548–L559, January 2010. doi: 10.1152/ajplung.00230.2010.

68. G. Bulut, S.-H. Hong, K. Chen, E. M. Beauchamp, S. Rahim, G. W. Kosturko, E. Glasgow, S. Dakshanamurthy, H.-S. Lee, I. Daar, J. A. Toretsky, C. Khanna, and A. Üren. Small molecule inhibitors of ezrin inhibit the invasive phenotype of osteosarcoma cells. Oncogene, 31(3):269–281, January 2012. doi: 10.1038/onc.2011.245.

69. James J. Tomasek, Melville B. Vaughan, Bradley P. Kropp, Giulio Gabbiani, Michael D. Martin, Carol J. Haaksma, and Boris Hinz. Contraction of myofibroblasts in granulation tissue is dependent on Rho/Rho kinase/myosin light chain phosphatase activity. Wound Repair and Regeneration, 14(3):313–320, 2006. doi: 10.1111/j.1743-6109.2006.00126.x.

70. James J. Tomasek, Giulio Gabbiani, Boris Hinz, Christine Chaponnier, and Robert A. Brown. Myofibroblasts and mechano-regulation of connective tissue remodelling. Nature Reviews Molecular Cell Biology, 3(5):349–363, May 2002. doi: 10.1038/nrm809.

71. Xiangwei Huang, Ying Gai, Naiheng Yang, Baogen Lu, Chrishan S. Samuel, Victor J. Thannickal, and Yong Zhou. Relaxin Regulates Myofibroblast Contractility and Protects against Lung Fibrosis. The American Journal of Pathology, 179(6):2751–2765, December 2011. doi: 10.1016/j.ajpath.2011.08.018.

72. Adam C. Midgley, Emma L. Woods, Robert H. Jenkins, Charlotte Brown, Usman Khalid, Rafael Chavez, Vincent Hascall, Robert Steadman, Aled O. Phillips, and Soma Meran. Hyaluronidase-2 Regulates RhoA Signaling, Myofibroblast Contractility, and Other Key Profibrotic Myofibroblast Functions. The American Journal of Pathology, 190(6):1236–1255, June 2020. doi: 10.1016/j.ajpath.2020.02.012.

73. Anne J. Ridley and Alan Hall. The small GTP-binding protein rho regulates the assembly of focal adhesions and actin stress fibers in response to growth factors. Cell, 70(3):389–399, August 1992. doi: 10.1016/0092-8674(92)90163-7.

74. M. Chrzanowska-Wodnicka and K. Burridge. Rho-stimulated contractility drives the formation of stress fibers and focal adhesions. Journal of Cell Biology, 133(6):1403–1415, June 1996. doi: 10.1083/jcb.133.6.1403.

75. Kiran Bhadriraju, Michael Yang, Sami Alom Ruiz, Dana Pirone, John Tan, and Christopher S. Chen. Activation of ROCK by RhoA is regulated by cell adhesion, shape, and cytoskeletal tension. Experimental Cell Research, 313(16):3616–3623, October 2007. doi: 10.1016/j.yexcr.2007.07.002.

76. Masayoshi Uehata, Toshimasa Ishizaki, Hiroyuki Satoh, Takashi Ono, Toshio Kawahara, Tamami Morishita, Hiroki Tamakawa, Keiji Yamagami, Jun Inui, Midori Maekawa, and Shuh Narumiya. Calcium sensitization of smooth muscle mediated by a Rho-associated protein kinase in hypertension. Nature, 389(6654):990–994, October 1997. doi: 10.1038/40187.

77. Mian Zhou and Yu-Li Wang. Distinct Pathways for the Early Recruitment of Myosin II and Actin to the Cytokinetic Furrow. Molecular Biology of the Cell, 19(1):318–326, October 2007. doi: 10.1091/mbc.e07-08-0783.

78. L. G. Cao and Y. L. Wang. Mechanism of the formation of contractile ring in dividing cultured animal cells. II. Cortical movement of microinjected actin filaments. Journal of Cell Biology, 111(5):1905–1911, November 1990. doi: 10.1083/jcb.111.5.1905.

79. A. Chen, P D. Arora, C. A. McCulloch, and A. Wilde. Cytokinesis requires localized β-actin filament production by an actin isoform specific nucleator. Nature Communications, 8(1): 1530, November 2017. doi: 10.1038/s41467-017-01231-x.

80. Sara O. Dean, Stephen L. Rogers, Nico Stuurman, Ronald D. Vale, and James A. Spudich. Distinct pathways control recruitment and maintenance of myosin II at the cleavage furrow during cytokinesis. Proceedings of the National Academy of Sciences, 102(38):13473–13478, September 2005. doi: 10.1073/pnas.0506810102.

81. Yitong Li and Keith Burridge. Cell-Cycle-Dependent Regulation of Cell Adhesions: Adhering to the Schedule. BioEssays, 41(1):1800165, 2019. doi: 10.1002/bies.201800165.

82. Benoit Vianay, Fabrice Senger, Simon Alamos, Maya Anjur-Dietrich, Elizabeth Bearce, Bevan Cheeseman, Lisa Lee, and Manuel Théry. Variation in traction forces during cell cycle progression. Biology of the Cell, 110(4):91–96, 2018. doi: 10.1111/boc.201800006.

83. Matthew C. Jones, Janet A. Askari, Jonathan D. Humphries, and Martin J. Humphries. Cell adhesion is regulated by CDK1 during the cell cycle. Journal of Cell Biology, 217(9): 3203–3218, September 2018. doi: 10.1083/jcb.201802088.

84. Meredith E. K. Calvert, Graham D. Wright, Fong Yew Leong, Keng-Hwee Chiam, Yinxiao Chen, Gregory Jedd, and Mohan K. Balasubramanian. Myosin concentration underlies cell size-dependent scalability of actomyosin ring constriction. Journal of Cell Biology, 195(5):799–813, November 2011. doi: 10.1083/jcb.201101055.

85. Ting Gang Chew, Junqi Huang, Saravanan Palani, Ruth Sommese, Anton Kamnev, Tomoyuki Hatano, Ying Gu, Snezhana Oliferenko, Sivaraj Sivaramakrishnan, and Mohan K. Balasubramanian. Actin turnover maintains actin filament homeostasis during cytokinetic ring contraction. Journal of Cell Biology, 216(9):2657–2667, September 2017. doi: 10.1083/jcb.201701104.

86. Inês Mendes Pinto, Boris Rubinstein, Andrei Kucharavy, Jay R. Unruh, and Rong Li. Actin Depolymerization Drives Actomyosin Ring Contraction during Budding Yeast Cytokinesis. DevelopmentalCell, 22(6):1247–1260, June 2012. doi: 10.1016/j.devcel.2012.04.015.

87. Dietmar B. Oelz, Boris Y. Rubinstein, and Alex Mogilner. A Combination of Actin Treadmilling and Cross-Linking Drives Contraction of Random Actomyosin Arrays. Biophysical Journal, 109(9):1818–1829, November 2015. doi: 10.1016/j.bpj.2015.09.013.

88. Ian A Darby, Betty Laverdet, Frédéric Bonté, and Alexis Desmoulière. Fibroblasts and myofibroblasts in wound healing. Clinical, Cosmetic and Investigational Dermatology, 7: 301–311, November 2014. doi: 10.2147/CCID.S50046.

89. Bin Li and James H-C. Wang. Fibroblasts and Myofibroblasts in Wound Healing: Force Generation and Measurement. Journal of tissue viability, 20(4):108–120, November 2011. doi: 10.1016/j.jtv.2009.11.004.

90. Keith Burridge and Erika S. Wittchen. The tension mounts: Stress fibers as force-generating mechanotransducers. Journal of Cell Biology, 200(1):9–19, January 2013. doi: 10.1083/jcb.201210090.

91. Kristopher E. Kubow and Alan Rick Horwitz. Reducing background fluorescence reveals adhesions in 3D matrices. Nature Cell Biology, 13(1):3–5, January 2011. doi: 10.1038/ncb0111-3.

92. Andrew D. Doyle, Nicole Carvajal, Albert Jin, Kazue Matsumoto, and Kenneth M. Yamada. Local 3D matrix microenvironment regulates cell migration through spatiotemporal dynamics of contractility-dependent adhesions. Nature Communications, 6(1):8720, November 2015. doi: 10.1038/ncomms9720.

93. Nicholas O. Deakin and Christopher E. Turner. Distinct roles for paxillin and Hic-5 in regulating breast cancer cell morphology, invasion, and metastasis. Molecular Biology of the Cell, 22(3):327–341, December 2010. doi: 10.1091/mbc.e10-09-0790.

94. Jill S. Harunaga and Kenneth M. Yamada. Cell-matrix adhesions in 3D. Matrix Biology, 30 (7):363–368, September 2011. doi: 10.1016/j.matbio.2011.06.001.

95. Stacey Lee and Sanjay Kumar. Actomyosin stress fiber mechanosensing in 2D and 3D. F1000Research, 5, September 2016. doi: 10.12688/f1000research.8800.1.

96. A. Seal, A. K. Dalui, M. Banerjee, A. K. Mukhopadhyay, and K. K. Phani. Mechanical properties of very thin cover slip glass disk. Bulletin of Materials Science, 24(2):151–155, April 2001. doi: 10.1007/BF02710092.

97. Aisling Ní Annaidh, Karine Bruyère, Michel Destrade, Michael D. Gilchrist, and Mélanie Otténio. Characterization of the anisotropic mechanical properties of excised human skin. Journal of the Mechanical Behavior of Biomedical Materials, 5(1):139–148, January 2012. doi: 10.1016/j.jmbbm.2011.08.016.

98. Marion Geerligs, Lambert van Breemen, Gerrit Peters, Paul Ackermans, Frank Baaijens, and Cees Oomens. In vitro indentation to determine the mechanical properties of epidermis. Journal of Biomechanics, 44(6):1176–1181, April 2011. doi: 10.1016/j.jbiomech.2011.01.015.

99. Nancy L. Halliday and James J. Tomasek. Mechanical Properties of the Extracellular Matrix Influence Fibronectin Fibril Assembly in Vitro. Experimental Cell Research, 217(1):109–117, March 1995. doi: 10.1006/excr.1995.1069.

100. Katsumi Mochitate, Pamala Pawelek, and Frederick Grinnell. Stress relaxation of contracted collagen gels: Disruption of actin filament bundles, release of cell surface fibronectin, and down-regulation of DNA and protein synthesis. Experimental Cell Research, 193(1):198–207, March 1991. doi: 10.1016/0014-4827(91)90556-A.

101. Robert J. Pelham and Yu-li Wang. High Resolution Detection of Mechanical Forces Exerted by Locomoting Fibroblasts on the Substrate. Molecular Biology of the Cell, 10(4):935–945, April 1999. doi: 10.1091/mbc.10.4.935.

102. Gabriele Nasello, Pilar Alamán-Díez, Jessica Schiavi, María Ángeles Pérez, Laoise McNamara, and José Manuel García-Aznar. Primary Human Osteoblasts Cultured in a 3D Microenvironment Create a Unique Representative Model of Their Differentiation Into Osteocytes. Frontiers in Bioengineering and Biotechnology, 8, 2020. doi: 10.3389/fbioe.2020.00336.

103. Wei Zhu, Nathan J. Castro, Haitao Cui, Xuan Zhou, Benchaa Boualam, Robert McGrane, Robert I. Glazer, and Lijie Grace Zhang. A 3D printed nano bone matrix for characterization of breast cancer cell and osteoblast interactions. Nanotechnology, 27(31):315103, June 2016. doi: 10.1088/0957-4484/27/31/315103.

104. R.-P Franke, M. Gräfe, H. Schnittler, D. Seiffge, C. Mittermayer, and D. Drenckhahn. Induc-tion of human vascular endothelial stress fibres by fluid shear stress. Nature, 307(5952): 648–649, February 1984. doi: 10.1038/307648a0.

105. A. J. Wong, T. D. Pollard, and I. M. Herman. Actin filament stress fibers in vascular endothelial cells in vivo. Science, 219(4586):867–869, February 1983. doi: 10.1126/science.6681677.

106. F. Grinnell. Fibroblasts, myofibroblasts, and wound contraction. Journal of Cell Biology, 124(4):401–404, February 1994. doi: 10.1083/jcb.124.4.401.

107. Penny K. Mar, Partha Roy, Helen L. Yin, H. Dwight Cavanagh, and James V Jester. Stress Fiber Formation is Required for Matrix Reorganization in a Corneal Myofibroblast Cell Line. Experimental Eye Research, 72(4):455–466, April 2001. doi: 10.1006/exer.2000.0967.

108. Nathan Sandbo, Andrew Lau, Jacob Kach, Caitlyn Ngam, Douglas Yau, and Nickolai O. Dulin. Delayed stress fiber formation mediates pulmonary myofibroblast differentiation in response to TGF-β. American Journal of Physiology-Lung Cellular and Molecular Physiology, 301(5):L656–L666, August 2011. doi: 10.1152/ajplung.00166.2011.

109. G. Gabbiani, B. J. Hirschel, G. B. Ryan, P. R. Statkov, and G. Majno. GRANULATION TISSUE AS A CONTRACTILE ORGAN A STUDY OF STRUCTURE AND FUNCTION. Journal of Experimental Medicine, 135(4):719–734, April 1972. doi: 10.1084/jem.135.4.719.

110. Charles R. Harris, K. Jarrod Millman, Stéfan J. van der Walt, Ralf Gommers, Pauli Virtanen, David Cournapeau, Eric Wieser, Julian Taylor, Sebastian Berg, Nathaniel J. Smith, Robert Kern, Matti Picus, Stephan Hoyer, Marten H. van Kerkwijk, Matthew Brett, Allan Haldane, Jaime Fernández del Río, Mark Wiebe, Pearu Peterson, Pierre Gérard-Marchant, Kevin Sheppard, Tyler Reddy, Warren Weckesser, Hameer Abbasi, Christoph Gohlke, and Travis E. Oliphant. Array programming with NumPy. Nature, 585(7825):357–362, September 2020. doi: 10.1038/s41586-020-2649-2.

111. Matplotlib: A 2D Graphics Environment - IEEE Journals & Magazine.

112. Pauli Virtanen, Ralf Gommers, Travis E. Oliphant, Matt Haberland, Tyler Reddy, David Cournapeau, Evgeni Burovski, Pearu Peterson, Warren Weckesser, Jonathan Bright, Stéfan J. van der Walt, Matthew Brett, Joshua Wilson, K. Jarrod Millman, Nikolay Mayorov, Andrew R. J. Nelson, Eric Jones, Robert Kern, Eric Larson, C. J. Carey, ìlhan Polat, Yu Feng, Eric W. Moore, Jake VanderPlas, Denis Laxalde, Josef Perktold, Robert Cim-rman, Ian Henriksen, E. A. Quintero, Charles R. Harris, Anne M. Archibald, Antônio H. Ribeiro, Fabian Pedregosa, and Paul van Mulbregt. SciPy 1.0: fundamental algorithms for scientific computing in Python. Nature Methods, 17(3):261–272, March 2020. doi: 10.1038/s41592-019-0686-2.

113. Boris Hinz, Vera Dugina, Christoph Ballestrem, Bernhard Wehrle-Haller, and Christine Chaponnier. α-Smooth Muscle Actin Is Crucial for Focal Adhesion Maturation in Myofibroblasts. Molecular Biology of the Cell, 14(6):2508–2519, February 2003. doi: 10.1091/mbc.e02-11-0729.

