## Supplemental Information for "Ventral Stress Fibers Induce Plasma Membrane Deformation in Human Fibroblasts"

### Supplementary Information

#### Fig. S1: Myofibroblast Validation.

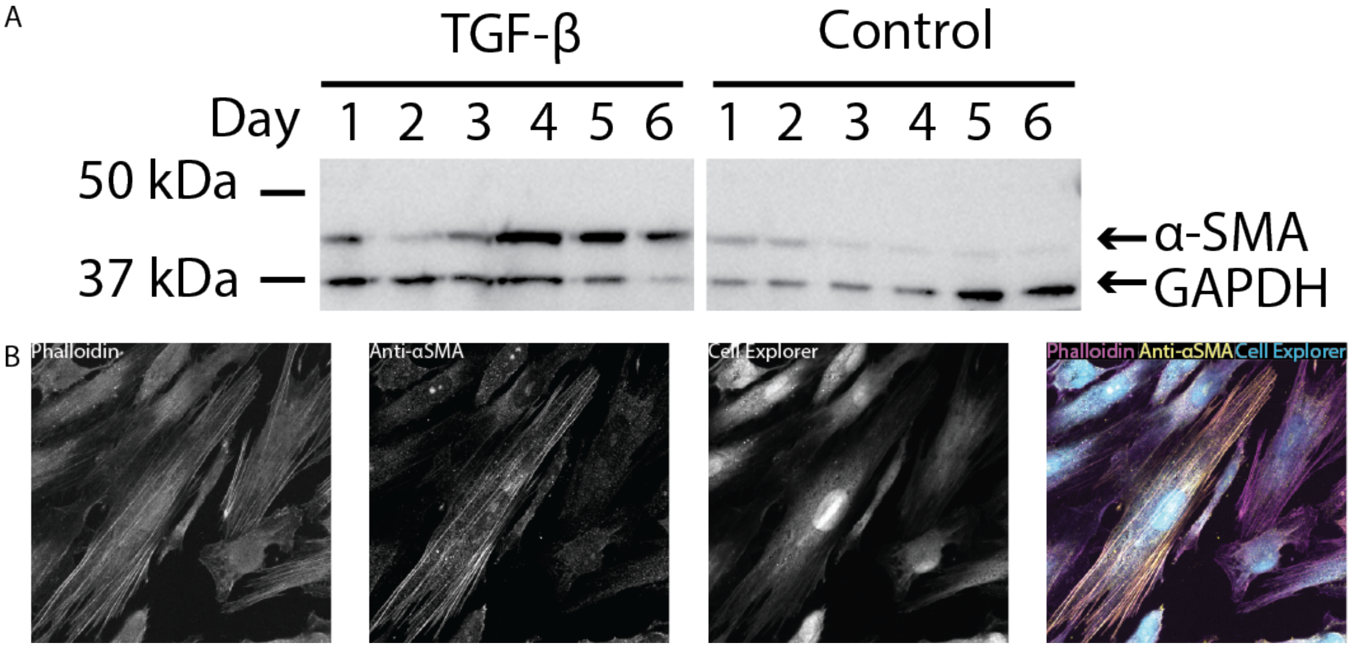


**Fig. S1. Validation of Fibroblast to Myofibroblast Transition** | **(A)** Cells were treated with TGF$\beta$-1 or control growth media for up to six days, with media replenishment every two days, and then a western blot was used to probe $\alpha$-SMA expression (anti-$\alpha$-SMA @ 1:500, anti-GAPDH @ 1:500), goat anti-mouse HRP secondary @ 1:1000). $\alpha$-SMA expression peaked after 4 days (96 hours) of treatment, which was the treatment time used for the rest of the study. **(B)** After 96 hours of TGF$\beta$-1 treatment, HDFs were loaded with Cell Explorer dye and then stained for $\alpha$-SMA and actin. Some cells showed colocalization of $\alpha$-SMA and actin stress fibers, while others did not. This is not unexpected as myofibroblasts come in two forms: proto-myofibroblasts that express $\alpha$-SMA without incorporating it into their cytoskeleton, and mature myofibroblasts that do incorporate it into their cytoskeleton (113). Given the destructive nature of staining for $\alpha$-SMA, both kinds of myofibroblasts were used in this study.

**Figure S2: TEM Images of Focal Adhesions.**

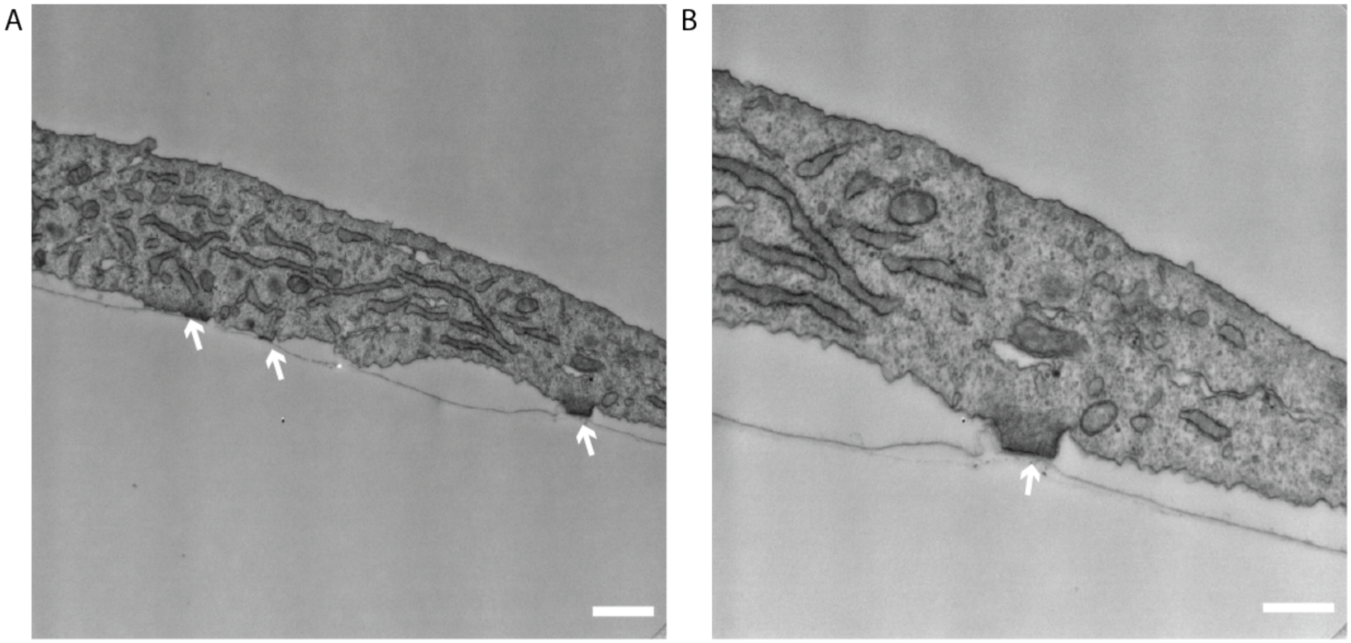


**Fig. S2. TEM images of Focal Adhesions | (A)** In these cross sections, clear focal adhesions can be seen (marked by white arrows). These focal adhesions are consistent with those observed by Abercrombie, et al. (27–29) having both a sharp edge boundary, as well as the distinctive dark region on the membrane-substrate interface, which gave rise to the initial observation of focal adhesions as electron-dense "plaques." The structures in Figure 3C have a much smoother curvature than the focal adhesions seen in these cells, and lack the electron dense plaque on the cell membrane. Scale Bar = 1 $\mu$m. **(B)** A zoomed-in region from the same cell as **(A)**, focusing on a single focal adhesion (white arrow). Scale Bar = 500 nm.

**Fig. S3: TEM Images of Stress Fiber-Induced Membrane Bending.**

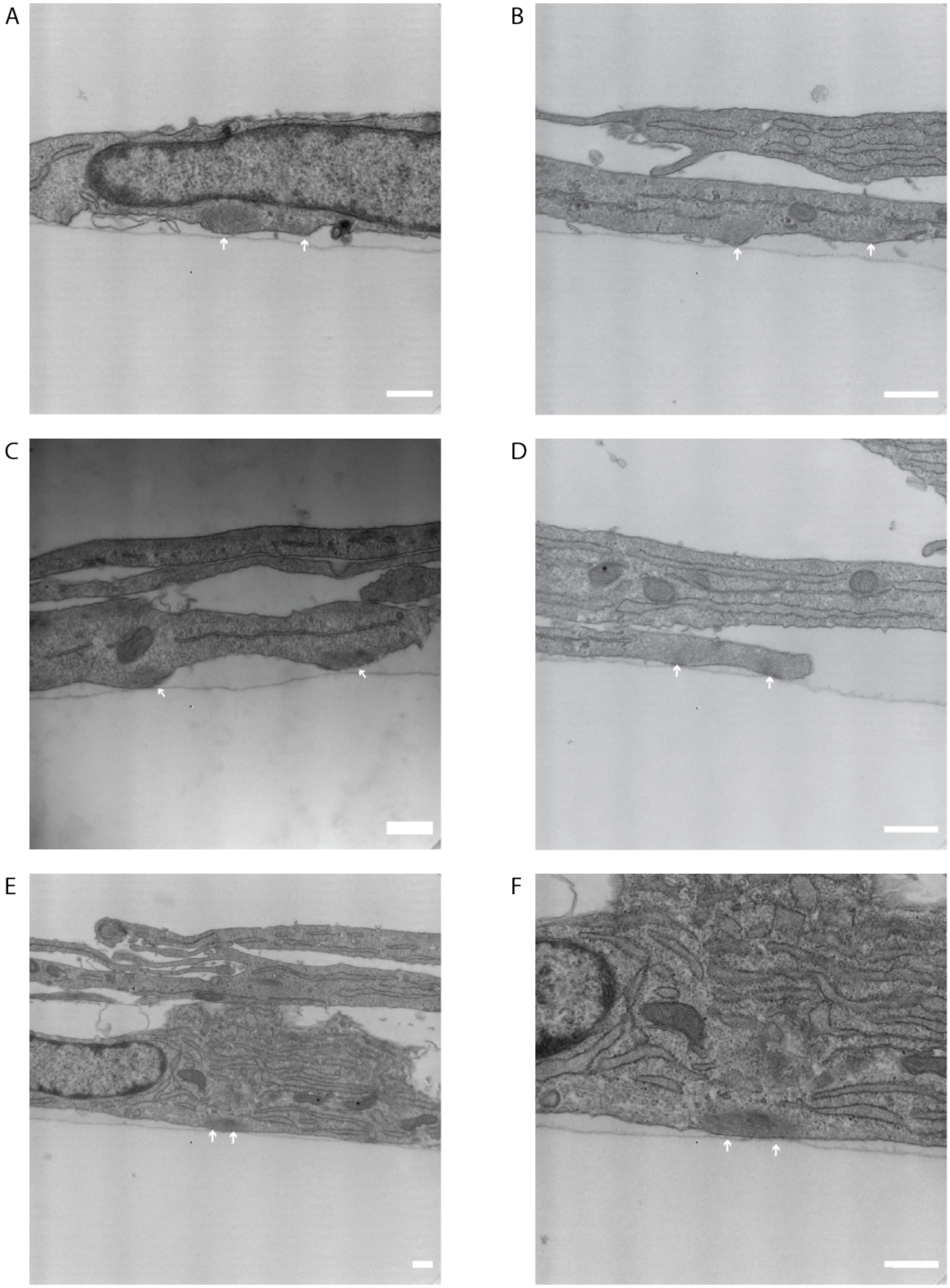


**Fig. S3. TEM images of Stress Fiber-Induced Membrane Bending | (A-E)** Images of stress fiber induced membrane bending, marked by white arrows. **(F)** is a zoomed in region of **(E)**. Scale Bar = 500 nm.
